# Single-cell RNA-Seq data have prevalent blood contamination but can be rescued by Originator, a computational tool separating single-cell RNA-Seq by genetic and contextual information

**DOI:** 10.1101/2024.04.04.588144

**Authors:** Thatchayut Unjitwattana, Qianhui Huang, Yiwen Yang, Leyang Tao, Youqi Yang, Mengtian Zhou, Yuheng Du, Lana X. Garmire

## Abstract

Single-cell RNA sequencing (scRNA-Seq) data from complex human tissues have prevalent blood cell contamination during the sample preparation process. They may also comprise cells of different genetic makeups. These issues demand rigorous preprocessing and filtering prior to the downstream functional analysis, to avoid biased conclusions due to cell types not of interest. Towards this, we propose a new computational framework, Originator, which deciphers single cells by the genetic origin and separates immune cells of blood contamination from those of expected tissue-resident cells. We demonstrate the accuracy of Originator at separating immune cells from the blood and tissue as well as cells of different genetic origins, using a variety of artificially mixed and real datasets. We show the prevalence of blood contamination in scRNA-Seq data of many tissue types. We alert the significant consequences if failing to adjust for these confounders, using scRNA-Seq data of pancreatic cancer and placentas as examples.

## Background

Single-cell RNA sequencing (scRNA-Seq) data analysis of complex (*e.g*. placenta) or inherently mixed (*e.g*. tumor) tissues pose significant challenges in computational biology. Many tissues contain blood vessels, and the resulting scRNA-seq data often include blood cell types (e.g., T cells) having highly similar expression profiles with the same cell types in the resident tissues. For example, in tumor studies precisely separating blood and tissue-resident immune cells in tumor tissues is crucial for understanding the tumor microenvironment [1]. In placenta tissues interfacing with the mother and fetus, isolating the maternal and fetal cells is pivotal to reveal cellular and immunological differences between them [2]. Therefore, a rigorous preprocessing step is required to enhance the quality of the data of interest, subsequently improving the downstream analysis. However, currently there is no dedicated tool to perform such a function. Here we fill in the void and propose a novel computational pipeline called Originator. It deconvolutes barcoded cells into different origins using inferred genotype information from scRNA-Seq data, as well as separating cells in the blood from those in solid tissues, an issue often encountered in scRNA-Seq experimentation.

## Results and Discussion

The proposed Originator framework is illustrated in **Fig 1a**. Cells undergo standard scRNA-Seq preprocessing steps, including quality control (QC), normalization, data integration, clustering, and cell-type annotation [3]. Then, Originator takes advantages of the differences in the genotype information from the scRNA-Seq reads, and utilize freemuxlet, a reference-free version of the popsicle suite [4] to separate the barcoded cells into different (default N=2) genetic origins, for example, maternal vs. fetal origins. Freemuxlet is chosen, as it showed better accuracies at recovering the genetic origins in comparison to scSplit [5], in the scRNA-Seq data of two placentas and the paired clear cell renal cell carcinoma (ccRCC) and PBMC cells from the same patient (**Supplementary Table 1**). Next, given that many tissues have blood contamination, the subsequent step separates immune cells by blood vs. expected tissue-resident context using the publicly available whole blood scRNA-Seq data as the reference [6]. After dissecting the heterogeneity of the scRNA-seq data, various downstream functional analysis can be performed.

**Fig. 1.**
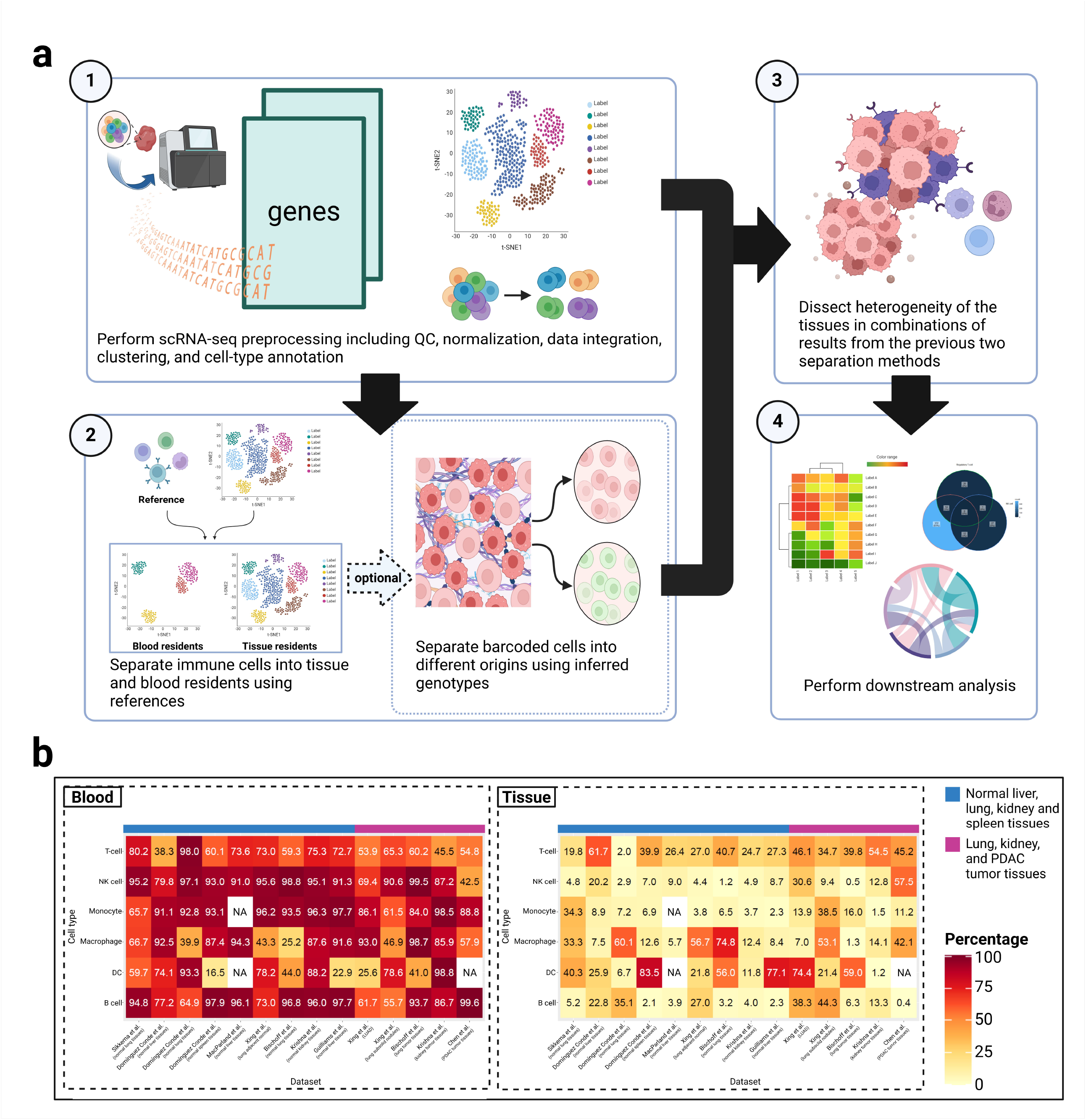
Illustration of Originator framework and the prevalence of blood immune cells in tissue samples. **a**, The input data are the scRNA-Seq experiment on tissue sections. (1) data preprocessing and cell type annotation. (2) separating barcoded cells into different origins by blood vs tissue residents context and optionally by inferred genotype information. (3) using the results in steps (1) and (2) to dissect tissue heterogeneity. (4) the functional downstream analyses with respect to cells’ origins. **b**, Heatmap plots show inferred immune cell type proportions originated from the blood (left) vs. the tissue (right), from scRNA-Seq data of a wide-variety of organs including liver, lung, spleen, kidney, pancreas and in the normal or cancer tissues. The cell types were annotated by the original publications and blood vs tissue identification were done by Originator. NA: the cell type does not exist in the original publication.

Current public scRNA-Seq data lack paired PBMC (or blood) and complex primary tissue completely free of blood contamination from the same person. We therefore first validated the pipeline using the artificially mixed data from a PBMCs dataset and a cell mixture dataset containing three breast cancer lines (T47D, BT474, MCF7), monocytes, lymphocytes, and stem cells **(Supplementary Notes)** [7–9]. Before creating the artificial mixture scRNA-Seq data, we removed the batch effect between the PBMC and the breast cancer cell lines (**Supplementary Figure 1)**, using Harmony [10] which showed good performance for this task previously [11]. As shown by the UMAP plot in **Supplementary Figure 1**, Originator separates the two compartments highly accurately, with AUC of 0.96, F-1 score of 0.97 and Area under the Precision-Recall Curve (AUCPR) of 0.96 (**Supplementary Table 2)**. Next, we tested Originator on the dataset of Krishna et al. 2021 [12], which contains paired ccRCC tissue and PBMC from the same patients. To generate the “ground truth” cell types for blood-eliminated ccRCC tissue, we applied Originator (or Seurat, for comparison) to the ccRCC to remove the potential blood immune cells from those residing in the tissue. We then integrated the cleaned ccRCC tissues and PBMC to generate an artificial mixture. Next, we ran Originator on this pre-cleaned mixture dataset for five iterations, and obtained averaged F-scores of 0.98, 0.93, 0.91, and 0.99 for B cell, CD4 T-cell, CD8 T-cell and NK cell respectively on the pre-cleaned, mixed dataset (**Supplementary Figure 2**). We thus conclude that Originator is effective at removing blood contamination from the results of both datasets above.

To signify the prevalence of blood immune cells in tissue samples, we applied Originator to eight publicly available scRNA-Seq data from a variety of organs including lung, liver, spleen, kidney, and pancreas, with normal (or adjacent normal) and/or tumor tissue samples [13–20]. As summarized in **Fig.1b**, in most datasets the majority of the immune cells are from the blood, rather than the tissue. The detailed side-by-side comparison of UMAP plots of blood vs tissue immune cells is in **Supplementary Figure 3-4**. These results unambiguously demonstrate the vast amount of immune cells from blood, rather than the tissue microenvironment. Failure to remove these immune cells may yield significantly biased results.

To directly demonstrate the potential biases in the downstream analyses due to contamination of the immune cells from the blood, we first applied Originator to a pancreatic ductal adenocarcinoma (PDAC) scRNA-seq dataset [20], to demonstrate its utility in removing cells that originate from blood. Originator successfully separates immune cells in the blood from those in the tumor tissue **(Fig 2a)**, and mast and Myeloid-derived suppressor cells (MDSCs) are exclusive in tissues as expected. The immune cell type proportions of the two compartments are shown in **Fig 2b**. The blood has larger proportions of macrophages and regulatory T-cells, but fewer T-cells. We then performed the differential expression (DE) analysis comparing the common immune cell types between the blood and those expected from the tumor tissues, and detected significant differences in gene expression in each immune cell type, especially for macrophage and T-cells **(Fig. 2c, Supplementary Table 3-6)** [21,22]. For example, CCL4 and CCL5 expression is higher in T-cells in the tumor tissue compared to those in the blood (**Supplementary Figure 5a**), consistent with the previous study that these genes are directly associated with T-cell-inflamed phenotype and antigen-presenting cell mediated processes in PDAC [23]. We also examined the genes common in some immune cell types, but show significant differences between the blood vs. tumor compartments (**Supplementary Notes**). We subsequently performed Gene Set Enrichment Analysis (GSEA) comparing the immune cells’ gene expression difference between the blood vs tissue compartments **(Fig. 2d)**. Toll-like receptor (TLR) signaling pathway and NOD-like receptor signaling pathway are more active in expected tissue-resident T-cells and macrophages compared to those in blood. However, natural killer cell-mediated cytotoxicity signaling pathway is downregulated in NK cells from the tumor compared to the blood compartment, consistent with observed NK cell dysfunction in PDAC via the reduced cytotoxic granule components, granzyme B, and perforin [24]. To demonstrate the impact of blood cell contamination in functional interpretation, we inferred cell-cell communications CCC in tumor tissues before and after removing blood cells **(Fig 2e,f)**.

**Fig. 2.**
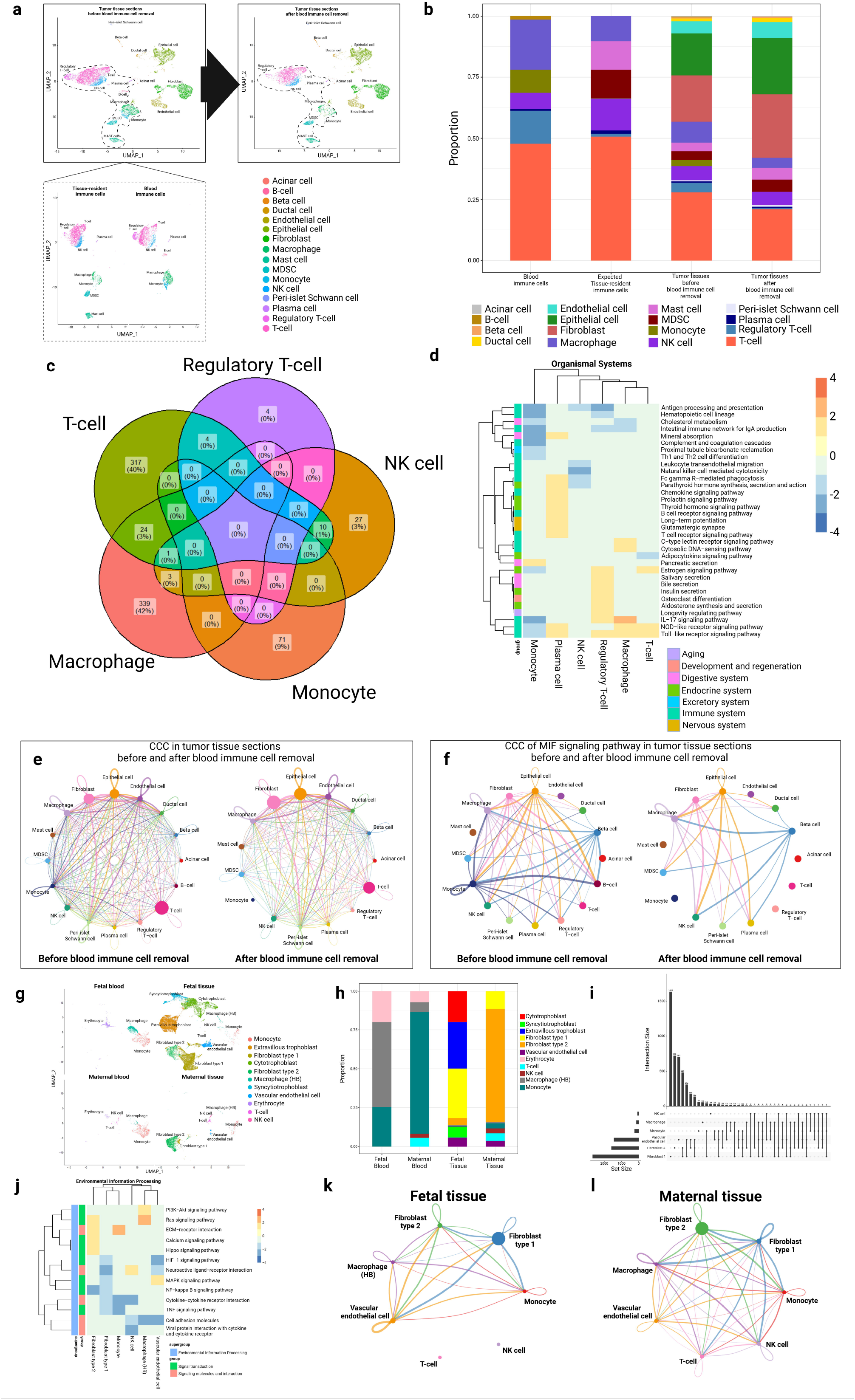
Applications of Originator to pancreatic ductal adenocarcinoma (PDAC) and placenta scRNA-seq data. **a**, UMAP plot of cells in PDAC tissues (top-left) separated into blood and expected tissue-resident immune cells (bottom-left). UMAP plot of cells in PDAC tissues after blood immune cell removal is shown on the top-right. **b**, The cell type proportion barplot shows the immune cell types in blood and PDAC tumor tissue, as well as all cell types in the tumor tissue before and after blood immune cell removal. **c**, Venn diagram of the significant DE genes identified when comparing immune cells in blood and expected tumor resident tissue. **d**, GSEA results comparing blood and expected tissue-resident immune cell types. **e**, Overall CCC in the tumor tissues before and after blood immune cell removal. **f**, CCC of MIF signaling pathway in tumor tissues before and after blood immune cell removal. **g**, UMAP plot of cells in placenta tissues deciphered into fetal vs. maternal origins and blood vs. placenta-resident cells. **h**, Cell type proportions in the different compartments related to the placenta. **i**, Upset plot of significant DE genes identified in common cell types between fetal and maternal placenta tissues. T-cell is excluded due to the near 0 counts in the fetal tissue. **j**, GSEA results comparing fetal and maternal tissues among common cell types. **k-l**, CCC of common cell types in fetal (**k**) and maternal (**l**) tissues.

While the interactions among the cell types are globally less frequent/noisy in the tumor compartment, those between fibroblast/epithelium and other cell types are strengthened in blood cells, as reflected by the increased node size of these two cell types **(Fig 2e)**. Such a trend of change is most drastic for the MIF signaling pathway (**Fig 2f**), which was previously reported to drive the malignant character of pancreatic cancer [25]. Particularly, MIF signaling between T-cells/regulatory T-cells and other cell types (eg. epithelial cells) are absent after removing blood immune cells, indicating a compromised anti-tumor immune response in PDAC [26].

Some tissues may have different genetic lineages that also need to be addressed. For example, the placenta has both maternal and fetal tissues. We then applied Originator to the human placenta [27], and separated the single cells into fetal and maternal origins as well as blood vs. expected tissue-resident cells. The cells from the fetal origin account for the majority of the cell populations, and the expected tissue-resident cells significantly over-weigh blood cells, as expected **(Fig 2g)**. The framework correctly and exclusively assigns trophoblast cells, including cytotrophoblasts, syncytiotrophoblasts, and extravillous trophoblasts, to the fetal tissue (**Fig 2g,2h**) [28]. On the contrary, immune cells, vascular endothelial cells, and fibroblasts (type 1 and type 2) appear in both maternal and fetal tissues, also as expected **(Fig 2h)**. We further performed DE analysis on the common cell types between fetal and maternal tissues and discovered drastic differences in gene expression related to the local tissue environment **(Fig 2i, Supplementary Table 7-9)**. For example, The top-ranked DE gene EGFL6 between maternal vs. fetal fibroblast cells is highly expressed in fibroblasts subtype 1 in fetal tissues as expected (**Supplementary Figure 6a**) [29]. However, it is much lower in fibroblast subtype 1 or mostly absent in fibroblast subtype 2 in maternal tissues, similar to a previous report [30]. Among the DE genes from macrophage cells (**Supplementary Table 9**), SEPP1 is expressed at much higher levels in Hofbauer cells originating from the fetal tissue compared to the maternal tissue (**Supplementary Figure 6b)**, in accordance with a previous study [31]. We additionally performed GSEA on the DE genes of the common cell types between fetal and maternal cells in the placenta tissue **(Fig 2j)**. The MAPK signaling pathway is enriched in vascular endothelial cells from the fetal tissues, which agrees with its role in growth factor-induced fetoplacental angiogenesis [32]. On the other hand, fibroblast type 2 from the fetal origin is enriched with several other signaling pathways, such as calcium, hippo, and Ras signaling pathways as well as ECM-receptor interaction **(Fig 2j)**, which indicates extracellular matrix (ECM) remodeling during trophoblast differentiation [33]. We also compared the CCC among the common cell types between the fetal and maternal tissues **(Fig 2k, 2l)**. Type 1 and type 2 fibroblasts both show different degrees of CCC with some other cell types when comparing the maternal and fetal tissue contexts. A higher interaction between fibroblast cells (type 1 and type 2) and vascular endothelial cells in fetal tissues reflects the active formation of the villous stroma underneath the syncytiotrophoblast and surrounding fetal capillaries [34]. On the contrary, higher interactions between type 2 fibroblast and macrophages in maternal tissue may assist trophoblast invasion through growth factors and cytokines [35]. Thus, teasing apart the cell types by their genetic and local tissue context helps to refine the molecular analysis in placenta scRNA-seq data.

## Discussion and conclusions

Originator is the first dedicated systematic tool to decipher scRNA-Seq data by genetic origin and blood/tissue contexts in heterogeneous tissues. It can be used as an effective tool to remove the undesirable blood cells in scRNA-Seq data. It can also provide improved cell type annotations and other downstream functional analyses, based on the genetic background.

Future work will be focused on generating pan-tissue immune cell atlas which are free of immune cells originating from blood contamination, which will better annotate the expected tissue-resident immune cells truthfully from the tissues of interest.

## Methods

### scRNA-Seq case study data sets

scRNA-Seq data from liver, lung, pancreas and kidney are used in this study. For lung, we used a large scRNA-Seq dataset from the integrated human lung cell atlas (v1.0), containing samples from 107 individuals [14], as well as lung cancer datasets from Xing et al. and Bischoff et al. [18,19]. For liver tissues, we used the single-cell liver landscape dataset from five individuals [16], and another healthy liver dataset from Guilliams et al. [17]. Additionally, we used lung, spleen, and liver scRNA-Seq datasets from Domínguez Conde et al. [15]. We obtained the paired blood and kidney cancer dataset from Krishna et al. [13]. For pancreatic ductal adenocarcinoma (PDAC), we use a scRNA-Seq dataset from GSE212966, which includes two tumor tissues [20]. The last dataset is the scRNA-Seq data from placenta tissues, comprising eight placenta samples from the previous study (EGA; https://www.ebi.ac.uk/ega/) hosted by the European Bioinformatics Institute (EBI; accession no. EGAS00001002449) [27]. Cell types in the placenta and PDAC tissues are annotated using the cell-type-specific marker genes [36](**Supplementary Table 10-11**).

### Description of Originator for scRNA-Seq data analysis

Originator is a multi-module framework that can be used to preprocess scRNA-Seq data from heterogeneous tissues. It consists of one mandatory step based on the tissue vs blood compartments and also an optional step based on the genetic origins. Originator takes the gene expression matrix processed by Cell Ranger (version 7.1.0) of 10x genomics as the input [37].

For genetic background based deciphering, it also uses the BAM file. It processes and outputs the R data serialization file (RDS). This output file contains gene expression and related information, including the cell type, blood and tissue immune cell annotation, and annotation by genetic information if this function is needed.

First, Originator separates immune cells in blood vs. those in the tissue using the blood immune cell scRNA-Seq reference constructed from the publicly available scRNA-seq data from the whole blood cells [6]. Annotated immune cells of interest (monocytes, macrophages, T-cells, regulatory T-cells, plasma cells, NK cells, and B-cells) from the datasets of this study are aligned to the whole blood scRNA-Seq reference data using package Seurat (4.3.0) [38]. For each immune cell type, the top 10 latent variables from each scRNA-Seq sample are used to compute a pairwise Euclidean distance matrix between each query immune cell and the reference whole blood. The latent variables can be obtained by UMAP or PCA based dimension reduction. We compared UMAP-based and PCA-based dimension reduction on the data of Krishna et al. [13], and found that the former yielded higher accuracies in detecting the immune cell types (**Supplementary Figure 7**). Thus we used UMAP-based dimension reduction in this report. To separate each query of immune cells into blood or expected tissue-resident cells, k-means clustering (default k = 2) is applied using Euclidean distances relative to the whole blood immune cell reference. The cluster more similar to the whole blood immune cell references is annotated as the blood immune cells, and the other cluster is determined as the expected tissue-resident immune cells.

If the tissue contains the mixture of cells of different genetic background, such as maternal and fetal origins, Originator provides an optional step to decipher the barcoded cells by the genetic origin relies on the overlapping genetic variants extracted from the scRNA-Seq data. Genetic variants are extracted by *bcftools* (version 1.17) [39]. Variants informative of genetic origins are determined by excluding single nucleotide polymorphisms (SNPs) with minor allele frequency (MAF) greater than x percentages (default x=10), using the 1000 Genomes Phase 3 reference panel (release date: 2013/05/02) [40]. The *freemuxlet* package is used to separate the single cells with the parameter ‘--nsample N’ (default N=2), as implemented in popscle (https://github.com/statgen/popscle) [4]. N denotes the number of genetic origins of the single cells (e.g., N = 2 for the maternal and fetal origins in a placenta scRNA-Seq data). Mosaic doublets (a single barcoded cell with a 50% genetic mixture from two individuals) are also identified in this step as part of the quality control. As trophoblast cells in the placenta are of fetal origin, the cluster that contains trophoblast cells is identified with the fetal origin, while the other cluster is marked as the maternal origin.

### Testing the performance of Originator

The artificially mixed tissue-blood-resident data were generated to assess the ability of Originator to separate blood cells and tissue immune residents. We included two datasets: (1) an artificially mixed data from a PBMCs dataset and a cell mixture dataset containing three breast cancer lines (T47D, BT474, MCF7), monocytes, lymphocytes, and stem cells [7–9]. (2) A paired pre-cleaned ccRCC tissues and PBMC from the same patient provided by Krishna et al. [13]. The details of the evaluation are in **Supplementary Notes**. Additionally, we also benchmarked freemuxlet [4](used in Originator) against scSplit [5] in identifying cells from different genetic origins using two different datasets, including (1) scRNA-Seq data of two placenta samples and (2) mixed PBMC scRNA-Seq data from two ccRCC patients provided by Krishna et al. [13,27] **(Supplementary Notes)**.

### Downstream analysis

DE analysis was performed using *FindMarkers()* in the package Seurat (4.3.0) [38]. GSEA is done using *gseKEGG()* in the package ClusterProfiler (4.8.2) with the gene set information from the Kyoto Encyclopedia of Genes and Genomes (KEGG) database [41,42]. CCC inference was performed using the package CellChat [43].

## Code availability

Originator is freely available to use at URL: https://github.com/lanagarmire/Originator

## Abbreviations

CCC: Cell-cell communication
DE: Differential expression
GSEA: Gene set enrichment analysis
PBMC: Peripheral blood mononuclear cells
PDAC: Pancreatic ductal adenocarcinoma
QC: Quality control
scRNA-Seq: Single-cell RNA sequencing

## Declarations

### Ethics approval and consent to participate

Not applicable

### Consent for publication

Not applicable

### Availability of data and materials

We use a healthy lung scRNA-Seq dataset from the integrated human lung cell atlas (v1.0), containing samples from 107 individuals [14]. Additionally, we use lung, spleen, and liver scRNA-Seq datasets from Domínguez Conde et al. [15]. For liver tissues, we used the single-cell liver landscape dataset from five individuals [16], as well as another healthy liver dataset from Guilliams et al. [17]. For cancer tissues, we incorporated lung cancer datasets from Xing et al. and Bischoff et al. [18,19], and a kidney cancer dataset from Krishna et al. [13]. For pancreatic ductal adenocarcinoma (PDAC), we use a scRNA-Seq dataset from GSE212966, which includes two tumor tissues [20]. The last dataset is the scRNA-Seq data from placenta tissues, comprising eight placenta samples from the previous study (EGA; https://www.ebi.ac.uk/ega/) hosted by the European Bioinformatics Institute (EBI; accession no. EGAS00001002449) [27]. For simulation, we generated the data using an 8k PBMCs dataset from a healthy donor (blood-resident cells) and an *in vitro* cell mixture (expected tissue-resident cells) containing three breast cancer lines (T47D, BT474, MCF7), monocytes (Thp1), lymphocytes (Jukrat), and stem cells (hMSC) [8,9].

### Authors’ contributions

L.X.G. and Q.H. conceived this project and supervised the study. T.U. modified the pipeline, carried the analysis, and wrote the manuscript. Q.H. wrote the Originator pipeline. M.Z., Y.Y., Y.Y., Y.D. and L.T. assisted the analysis and tested the package.

## Acknowledgements

The authors acknowledge all lab members of Garmire Group for helpful discussions.

