## Supplementary Information for "Single-cell RNA-Seq data have prevalent blood contamination but can be rescued by Originator, a computational tool separating single-cell RNA-Seq by genetic and contextual information"

**Supplementary Notes**

**Testing the performance of Originator by an artificially mixed PBMC and breast cancer cell lines**

To evaluate the performance of the pipeline, we tested the tissue-blood deconvolution on artificially mixed tissue-blood resident data. This data were generated by combining an 8k PBMCs dataset from a healthy donor (blood-resident cells) and an *in vitro* cell mixture (expected tissue-resident cells) containing three breast cancer lines (T47D, BT474, MCF7), monocytes (Thp1), lymphocytes (Jukrat), and stem cells (hMSC) [1,2] **(Supplementary Figure 1a-b)**. The result shows that the pipeline correctly separates 98.31% of cells (12,809 out of 13,029 cells) into blood and expected tissue-resident cells **(Supplementary Figure 1c-d, Supplementary Table 2)**.

**Testing the performance of Originator by a paired clear cell renal cell carcinoma (ccRCC) tissue and PBMC from the same patients**

To evaluate the performance of Originator in separating blood and expected tissue-resident immune cells in a more realistic dataset with minimal batch effect, we used the dataset of Krishna et al. [3] which contains paired ccRCC tissue and PBMC for the same patients. To retrieve the ground truth for ccRCC tissues without blood cells, we applied Originator on ccRCC tissues to remove potential blood cells to generate cleaned ccRCC tissues. These cleaned ccRCC tissues were subsequently combined with the paired PBMC to generate a mixture of blood and tissue-resident immune cells. We ran Originator on this artificial mixture of blood and tissue-resident immune cells for five iterations and calculated the F1 scores. Originator achieves F1 scores of 0.98, 0.93, 0.91, and 0.99 for B cell, CD4 T-cell, CD8 T-cell, and NK cell, respectively **(Supplementary Figure 2)**.

**Benchmarking freemuxlet (in Originator) with scSplit , on separating single cells by genetic origins.**

We chose freemuxlet to separate single cells by genetic origins in Originator, as it does not require known references for SNPs in the samples. Its performance was benchmarked with another reference-free tool scSplit [4] on two different datasets, one from our own placenta data, and another with the mixed PBMC data from two ccRCC patients [3]. The mixed PBMC dataset contains transcripts of 200 cells from each of two different individuals, with a total of 400 cells. Single Nucleotide Variants (SNV) were called with Freebayes [5] and fed into the scSplit with the bam file. The results in both datasets indicate that freemuxlet outperforms the scSplit in assigning cells back to their patient origin **(Supplementary Table 1)**.

**Other DE genes between blood and tissue for the PDAC dataset**

Besides significant DE genes in T-cells due to the microenvironment, macrophages have the most abundant DE genes between tumor and blood. For example, higher expression of INHBA is present in expected tissue-resident compared to blood macrophages **(Supplementary Figure 5b)**, consistent with the previously reported upregulation of INHBA by M2-phenotype macrophages in pancreatic cancer **(Supplementary Table 5)** [6]. We observed higher expression of GZMK in expected tissue-resident NK cells compared to those in blood, consistent with the previously reported expression of cytotoxicity signatures comparing blood and tumor NK cell populations **(Supplementary Figure 5c, Supplementary Table 4)** [7]. We also observed higher expression of LILRA5 in expected tissue-resident monocytes than those from blood **(Supplementary Figure 5d, Supplementary Table 6)**. This is consistent with the previous study, which found that the cross-linking of LILRA5 on monocytes induces the production of pro-inflammatory cytokines, suggesting the inflammatory response associated with PDAC [8].

Some DE genes between expected tissue-resident and blood immune cells are common among different immune cell types (**Supplementary Figure 8**). We observed that MT-ND1, TNFRSF4, RPS26, and LTB are differentially expressed in both T-cell and T-reg in PDAC tissues compared to those in blood. MT-ND1 has higher expression in the tissue compartment compared to the blood, consistent with the previous study [9]. On the other hand, the expression of TNFRSF4, RPS26, and LTB is decreased in T-cell and T-reg in PDAC tissues compared to those in blood, with RPS26 showing higher expression levels than the other two genes. MT-ND1 is crucial for ATP production, and its high expression in tissue-resident T-cells and T-reg may reflect elevated energy demands necessary for effective immune responses against tumors [10]. In T-reg, mtDNA, including MT-ND1, was shown to be increased in tumor tissue due to mitochondrial abnormalities to drive cGAS-STING signaling [11]. RPS26 is a ribosomal protein-encoding gene and plays a key role in regulating their survival [12]. The decreased RPS26 expression in the T-cells and T-regs in the tumor tissue suggests the survival impairment of both cell types in PDAC.

We observed that RGCC, CD63, and LGALS1 are differentially expressed in both macrophages in NK cells. RGCC has higher expression in tissue-resident macrophages in PDAC tissues, consistent with the previous study that RGCC is highly expressed in M2-macrophages in tumor tissues [13,14]. It is also highly expressed in tissue-resident NK cells compared in the blood, consistent with the previous study [15]. Conversely, CD63 and LGALS1 are decreased in macrophages and NK cells in tissues compared to the blood. CD63 is an M2 macrophage marker in PDAC [16]. Reduced CD63 may impair exosome production and secretion in the macrophages, leading to diminished intercellular communication necessary for immune activation within the TME [17,18]. CD63 is expressed when NK cells are activated and ready to release cytotoxic granules; decreased expression of CD63 in NK cells may suggest impaired NK cell function within PDAC TME [18,19]. LGALS1 was previously shown to regulate tumor infiltration of macrophages [20]. Lower LGALS1 levels in tissue macrophages might reduce its infiltration, potentially impacting stromal remodeling [21,22].

ZNF331 is differentially expressed in NK cell, T-cell, and macrophage in the dataset. In NK cells, ZNF331 is higher in tissue-resident NK cells compared to those in blood. This is consistent with the expression of ZNF331 in tissue-resident NK cells in the previous studies [23,24]. However, ZNF331 is decreased in T-cells and macrophages in tissues compared to those in blood. Upregulation of ZNF331 expression in tissue-resident T-cells was shown to dictate its cell cycle regulation [25]. Decreased ZNF331 expression in tissue-resident T cells in PDAC tissue may impair the ability to proliferate and function effectively, and lead to compromised anti-tumor immunity. Though ZNF331 expression on macrophages is not extensively studied, Zinc finger protein was previously shown to play a role in macrophage polarization [26]. Deregulated ZNF331 in tissue-resident macrophages may impair the polarization of macrophages and the anti-tumor function.

26.	Xiao F, Shen J, Zhou L, Fang Z, Weng Y, Zhang C, et al. ZNF395 facilitates macrophage polarization and impacts the prognosis of glioma.

**Supplementary Figures**

**Supplementary Figure 1: Originator accurately separates artificially mixed blood cells from expected tissue-resident cells. a,** Data generation. **b,** UMAP of artificially mixed blood and tissue-resident data grouped by cell types. **c,** UMAP of artificially mixed blood-tissue resident data grouped by the data sources. **d,** UMAP of blood immune and expected tissue-resident cells recovered by Originator. **e-f,** UMAP of the alignment of whole blood reference, and artificially mixed blood-tissue resident data after batch correction using Harmony grouped by data sources (e) and cell types (f)

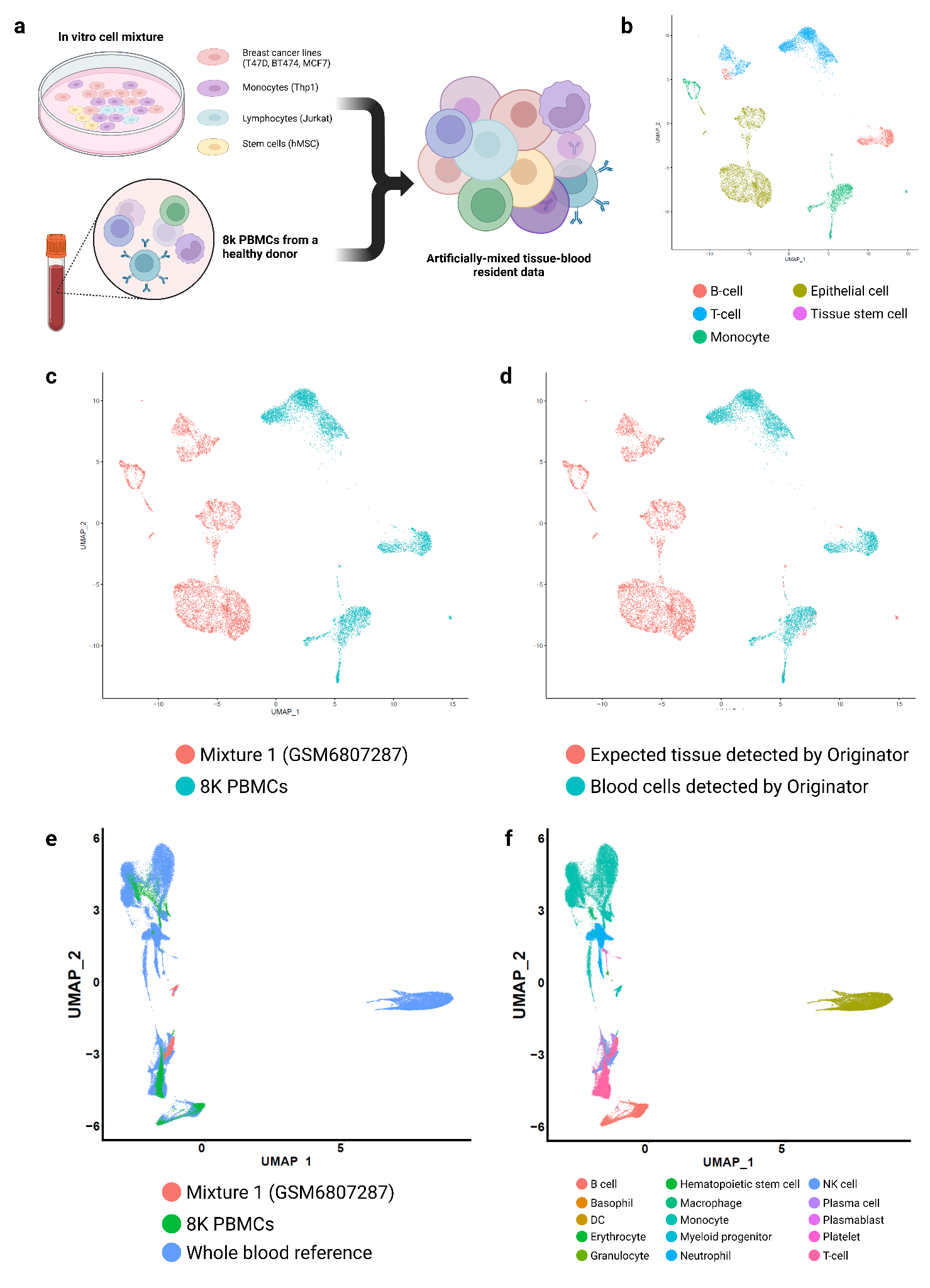

**Supplementary Figure 2: Originator accurately separates immune cells in the blood from those expected in the tissue of the same patient.** To generate the “ground truth” cell types for blood-eliminated ccRCC tissue, we applied Originator (or Seurat, for comparison) to ccRCC to remove the potential blood immune cells from ccRCC tissue first. We then integrated the cleaned ccRCC tissues and PBMC to generate an artificial mixture. Next we ran Originator on this pre-cleaned mixture dataset for five iterations.The boxplot shows the average F1 scores of separating immune cell types in the blood vs. blood-eliminated ccRCC tissue using Originator, where blood-elimination in the ccRCC data were done by Originator-based and Seurat-based approach, as stated above.

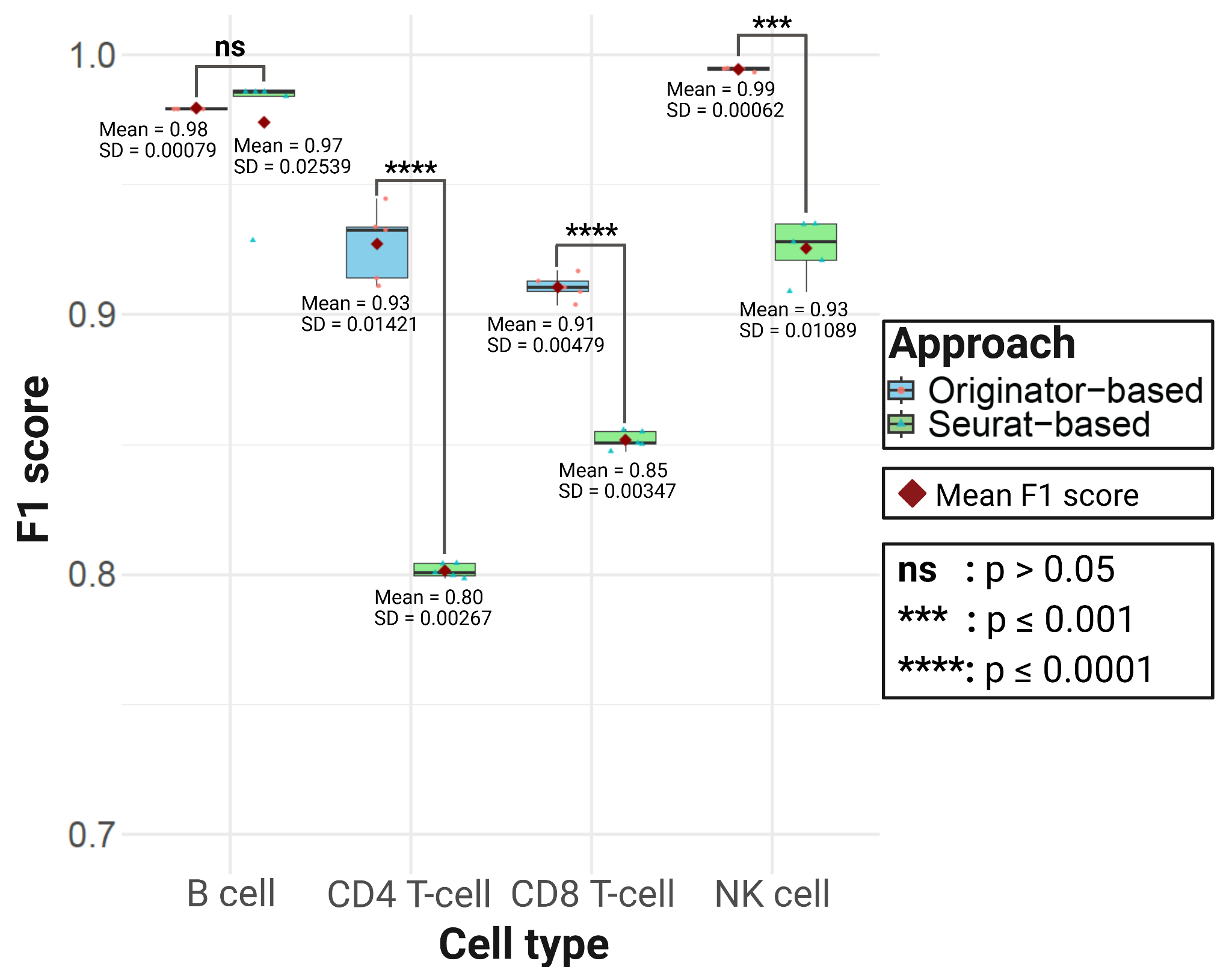

**Supplementary Figure 3:** **Blood and expected tissue-resident immune cells in healthy tissue datasets identified by Originator.** **a-b,** lung tissue datasets. **c,** spleen tissue datasets. **d-f,** liver tissue datasets.

**Supplementary Figure 4:** **Blood and expected tissue-resident immune cells in cancer tissue datasets identified by Originator. a-b,** lung cancer tissue datasets. **C,** kidney cancer tissue dataset.

**Supplementary Figure 5: UMAP displaying the expression of DE genes in immune cells from blood vs. expected PDAC tumor-resident immune cells. a,** CCL4 and CCL5 expression in T-cells. **b,** INHBA expression in macrophages. **c,** GZMK expression in NK cells. **d),** LILRA5 expression in monocytes.

**Supplementary Figure 6: Expression of DE genes in common cell types between fetal and maternal tissues of placenta. a,** EGFL6 expression in fibroblast type 1 and 2. **b,** SEPP1 expression in macrophages (HB)

**Supplementary Figure 7: Comparison of PCA-based and UMAP-based Originator on assigning cells from paired ccRCC tissues and PBMC data provided by Krishna et al. 2021**

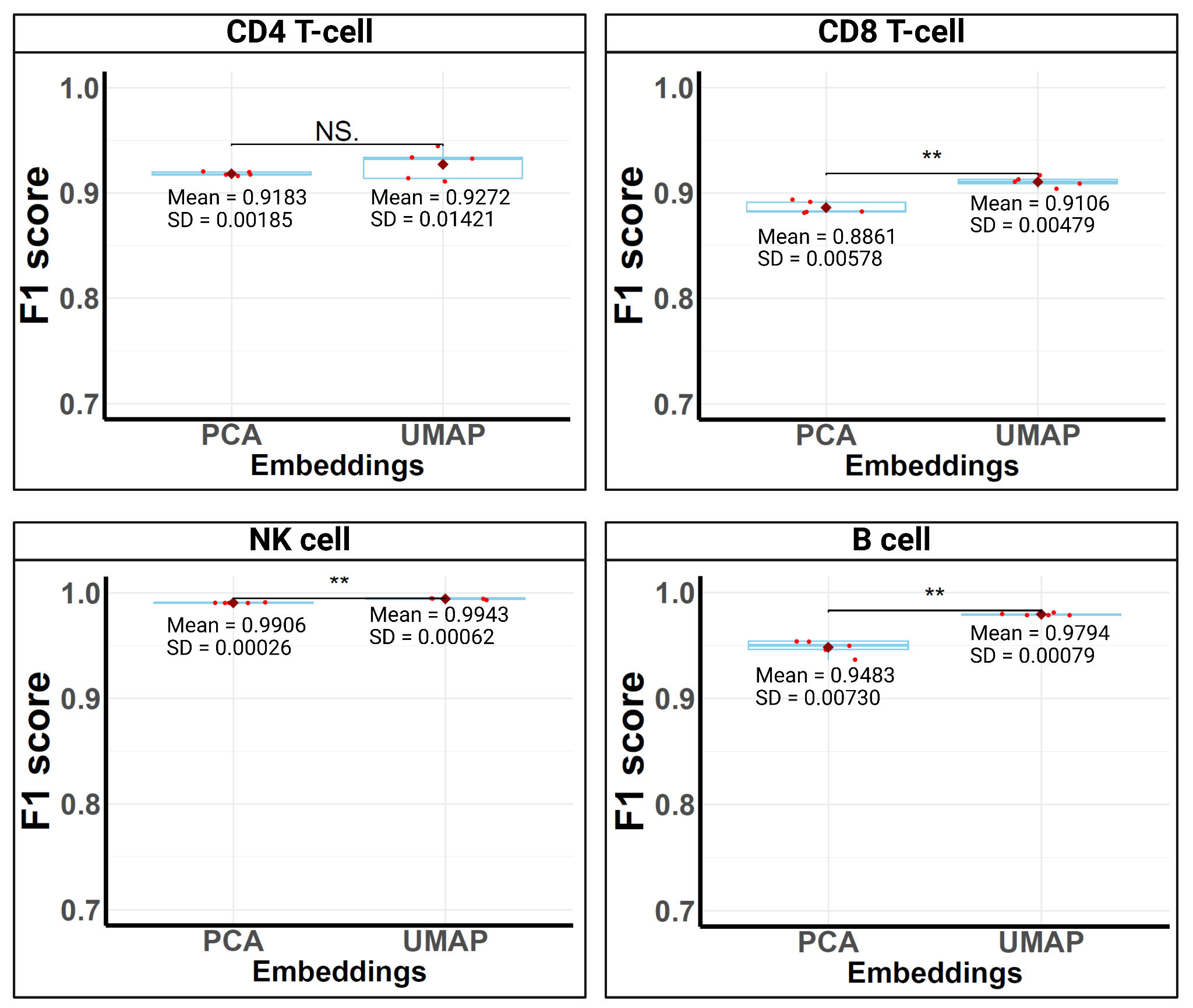

****: P ≤ 0.01**

**Supplementary Figure 8: UMAP plots showing the expression of DE genes between blood vs tissue among some common immune cell types;** A, UMAP of blood and tissue-resident immune cell., (b-e): DE genes common between T-cells and T-regs. (b) MT-ND1 (c) TNFRSF4 (d) RPS26 (e) LTB (f-h): DE genes common between macrophage and NK cells (f) RGCC (g) CD63. (h) LGALS1; (i): ZNF331,common in T-cell, NK cell, and macrophage cells

**Supplementary Table Legend**

**Supplementary Table 1:** Comparison between Freemuxlet and scSplit in assigning cells according to their genetic origins. Supplementary Table 1a-b show the performance of Freemuxlet and scSplit in (a) assigning trophoblast cells to the fetal origin in the placenta and (b) assigning cells according to genetic origin on mixed PBMC data from two clear cell renal cell carcinoma patients.

**Supplementary Table 2:** Average performance of 5 executions of blood and expected tissue-resident immune cell identification on the artificially-mixed blood-tissue resident data measured by four metrics including area under receiver operating characteristic curve (AUC), area under precision-recall curve (AUCPR), F1 score, and Matthews correlation coefficient (MCC).

**Supplementary Table 3:** Top significant DE genes comparing expected tissue-resident and blood T-cells in PDAC tumor tissue, as identified by Originator. log2 fold change (avg_log2FC) is averaged expression in expected tissue-resident compared to blood T-cells.

**Supplementary Table 4:** Top significant DE genes comparing expected tissue-resident and blood NK cells in PDAC tumor tissue, as identified by Originator. log2 fold change (avg_log2FC) is averaged expression in expected tissue-resident compared to blood NK cells.

**Supplementary Table 5:** Top significant DE genes comparing expected tissue-resident and blood macrophage cells in PDAC tumor tissue, as identified by Originator. log2 fold change (avg_log2FC) is averaged expression in expected tissue-resident compared to blood macrophage cells.

**Supplementary Table 6:** Top significant DE genes comparing expected tissue-resident and blood monocytes in PDAC tumor tissue, as identified by Originator. log2 fold change (avg_log2FC) is averaged expression in expected tissue-resident compared to blood monocytes.

**Supplementary Table 7:**Top significant DE genes comparing fetal and maternal fibroblast type 1 cells in placenta tissue, as identified by Originator. log2 fold change (avg_log2FC) is averaged expression in fetal compared to maternal fibroblast type 1.

**Supplementary Table 8:** Top significant DE genes comparing fetal and maternal fibroblast type 2 cells in placenta tissue, as identified by Originator. log2 fold change (avg_log2FC) is averaged expression in fetal compared to maternal fibroblast type 2.

**Supplementary Table 9:** Top significant DE genes comparing fetal and maternal macrophages in placenta tissue, as identified by Originator. log2 fold change (avg_log2FC) is averaged expression in fetal compared to maternal macrophages.

**Supplementary Table 10:** Cell-type-specific marker genes for placenta tissues

**Supplementary Table 11:** Cell-type-specific marker genes for PDAC tissues

**Supplementary Table 12:** Biological interpretation of exclusive genes between 1) T-cell and T-reg, 2) macrophage and NK cell, and 3) NK cell, T-cell, and macrophage
