## Supplementary Table for "Single-cell RNA-Seq data have prevalent blood contamination but can be rescued by Originator, a computational tool separating single-cell RNA-Seq by genetic and contextual information"

**Supplementary Tables**

**Supplementary Table 1: Comparison between Freemuxlet and scSplit in assigning cells according to their genetic origins.** Supplementary Table 1a-b show the performance of Freemuxlet and scSplit in (a) assigning trophoblast cells to the fetal origin and (b) assigning cells according to genetic origin on two individuals.

Supplementary Table 1a

| **Placenta tissue samples** | **Tools** | **Correct trophoblast assignment to the fetal origin (%)** |
| --- | --- | --- |
| Placenta sample 1 | Freemuxlet | 100 |
|  | scSplit | 52.24 |
| Placenta sample 2 | Freemuxlet | 100 |
|  | scSplit | 61.92 |

Supplementary Table 1b

| **Tools** | **F1 score** |
| --- | --- |
| Freemuxlet | 0.97 |
| scSplit | 0.73 |

**Supplementary Table 2: Average performance of 5 executions of blood and expected tissue-resident immune cell identification on the artificially-mixed blood-tissue resident data measured by four metrics including area under receiver operating characteristic curve (AUC), area under precision-recall curve (AUCPR), F1 score, and Matthews correlation coefficient (MCC).**

| **Cell type** | **AUC** | **AUCPR** | **F1 score** | **MCC** |
| --- | --- | --- | --- | --- |
| **T-cell** | 0.960  (SD = 0.001033) | 0.970  (SD = 0.000654) | 0.980  (SD = 0.000448) | 0.940  (SD = 0.001345) |
| **B-cell** | 0.998  (SD = 0) | 0.996  (SD = 0) | 0.997  (SD = 0) | 0.988  (SD = 0) |
| **monocyte** | 0.939  (SD = 0.007768) | 0.913  (SD = 0.007877) | 0.939  (SD = 0.005704) | 0.830  (SD = 0.015872) |

| **Overall performance** | | | |
| --- | --- | --- | --- |
| **AUC** | **AUCPR** | **F1 score** | **MCC** |
| 0.958  (SD = 0.002483) | 0.957  (SD = 0.002472) | 0.971  (SD = 0.001714) | 0.909  (SD = 0.005270) |

**Supplementary Table 3: Top significant DE genes comparing expected tissue-resident and blood T-cells in PDAC tumor tissue, as identified by Originator. log2 fold change (avg_log2FC) is averaged expression in expected tissue-resident compared to blood T-cells.**

|  | **p_val** | **avg_log2FC** | **pct.1** | **pct.2** | **p_val_adj** |
| --- | --- | --- | --- | --- | --- |
| CCL4 | 2.12E-36 | 1.573474 | 0.277 | 0.118 | 6.95E-32 |
| GZMK | 2.37E-60 | 1.333029 | 0.377 | 0.144 | 7.76E-56 |
| CCL5 | 2.94E-75 | 1.284637 | 0.507 | 0.222 | 9.62E-71 |
| PARP8 | 3.59E-60 | 1.284428 | 0.604 | 0.446 | 1.17E-55 |
| AOAH | 2.10E-14 | 1.034958 | 0.201 | 0.116 | 6.87E-10 |
| ZEB2 | 1.78E-21 | 1.004127 | 0.335 | 0.222 | 5.82E-17 |
| CBLB | 6.15E-22 | 0.952122 | 0.496 | 0.436 | 2.01E-17 |
| FOSB | 1.16E-21 | 0.868436 | 0.477 | 0.398 | 3.81E-17 |
| FKBP5 | 4.16E-14 | 0.808336 | 0.387 | 0.322 | 1.36E-09 |
| NKG7 | 6.24E-23 | 0.802629 | 0.18 | 0.071 | 2.04E-18 |
| GZMA | 3.41E-21 | 0.802536 | 0.311 | 0.193 | 1.12E-16 |
| ID2 | 2.56E-09 | 0.792395 | 0.291 | 0.237 | 8.38E-05 |
| METRNL | 3.48E-13 | 0.788526 | 0.234 | 0.155 | 1.14E-08 |
| ARHGAP26 | 8.32E-13 | 0.752598 | 0.353 | 0.288 | 2.72E-08 |
| CST7 | 1.02E-09 | 0.752057 | 0.218 | 0.156 | 3.34E-05 |
| TSC22D3 | 4.26E-20 | 0.734774 | 0.596 | 0.58 | 1.40E-15 |
| MT-ND1 | 7.77E-104 | 0.730171 | 0.98 | 0.989 | 2.54E-99 |
| NFE2L3 | 1.12E-11 | 0.72918 | 0.14 | 0.074 | 3.65E-07 |
| ZSWIM4 | 8.14E-09 | 0.711702 | 0.127 | 0.072 | 0.000266 |
| MT-CO1 | 2.42E-131 | 0.708822 | 0.995 | 0.997 | 7.93E-127 |
| CD8A | 8.88E-15 | 0.708242 | 0.109 | 0.041 | 2.91E-10 |
| AUTS2 | 1.51E-10 | 0.690459 | 0.247 | 0.175 | 4.96E-06 |
| PLCB1 | 1.24E-11 | 0.68456 | 0.279 | 0.199 | 4.04E-07 |
| MT-CO3 | 6.79E-116 | 0.680299 | 0.992 | 0.997 | 2.22E-111 |
| SLC7A5 | 1.18E-08 | 0.659229 | 0.297 | 0.251 | 0.000386 |
| SLC35F1 | 8.71E-09 | 0.640933 | 0.245 | 0.187 | 0.000285 |
| MT-ATP6 | 1.13E-116 | 0.637472 | 0.998 | 1 | 3.71E-112 |
| MT-ND5 | 1.34E-34 | 0.621219 | 0.781 | 0.814 | 4.39E-30 |
| RALGAPA1 | 5.56E-10 | 0.60728 | 0.54 | 0.564 | 1.82E-05 |
| MT-ND2 | 1.46E-55 | 0.597805 | 0.961 | 0.976 | 4.77E-51 |
| MT-CYB | 1.28E-68 | 0.574227 | 0.989 | 0.992 | 4.20E-64 |
| MT-CO2 | 1.70E-95 | 0.560024 | 0.995 | 0.997 | 5.57E-91 |
| CTSW | 1.02E-08 | 0.503019 | 0.108 | 0.057 | 0.000333 |
| MT-ND4 | 5.17E-57 | 0.494535 | 0.977 | 0.994 | 1.69E-52 |
| MT-ND3 | 1.43E-47 | 0.462944 | 0.978 | 0.99 | 4.68E-43 |
| SIK3 | 1.29E-07 | 0.436137 | 0.695 | 0.792 | 0.004238 |
| FOS | 2.23E-07 | 0.309387 | 0.554 | 0.517 | 0.007291 |
| SUMO2 | 2.00E-24 | -0.25005 | 0.238 | 0.46 | 6.53E-20 |
| SERF2 | 4.47E-20 | -0.25007 | 0.482 | 0.737 | 1.46E-15 |
| RPS19 | 1.02E-21 | -0.25049 | 0.928 | 0.976 | 3.35E-17 |
| BTG2 | 1.20E-20 | -0.25189 | 0.305 | 0.522 | 3.93E-16 |
| ERO1L | 5.18E-20 | -0.25264 | 0.065 | 0.169 | 1.70E-15 |
| CCM2 | 5.45E-22 | -0.2527 | 0.095 | 0.224 | 1.78E-17 |
| OGDH | 4.05E-24 | -0.2536 | 0.189 | 0.382 | 1.33E-19 |
| TTC39C | 6.61E-26 | -0.25397 | 0.284 | 0.519 | 2.17E-21 |
| EIF3F | 2.11E-26 | -0.25411 | 0.179 | 0.379 | 6.90E-22 |
| CD69 | 1.55E-11 | -0.25552 | 0.343 | 0.523 | 5.07E-07 |
| COMMD6 | 1.46E-27 | -0.25558 | 0.279 | 0.528 | 4.78E-23 |
| RPL38 | 7.96E-19 | -0.25615 | 0.715 | 0.894 | 2.61E-14 |
| RPL27A | 6.41E-24 | -0.25647 | 0.86 | 0.967 | 2.10E-19 |
| KRT10 | 9.37E-23 | -0.25653 | 0.111 | 0.255 | 3.07E-18 |
| TRAF3 | 3.02E-26 | -0.25657 | 0.179 | 0.376 | 9.90E-22 |
| TET2 | 2.32E-20 | -0.25666 | 0.11 | 0.242 | 7.60E-16 |
| CTSB | 2.13E-22 | -0.25709 | 0.06 | 0.169 | 6.98E-18 |
| TNFSF8 | 1.36E-17 | -0.2571 | 0.072 | 0.172 | 4.44E-13 |
| RPL36A | 1.56E-22 | -0.25711 | 0.398 | 0.653 | 5.11E-18 |
| ZNF706 | 5.81E-24 | -0.25727 | 0.085 | 0.217 | 1.90E-19 |
| MORC3 | 4.50E-24 | -0.25917 | 0.106 | 0.25 | 1.47E-19 |
| H3F3A | 5.01E-22 | -0.2592 | 0.527 | 0.783 | 1.64E-17 |
| NR3C1 | 1.54E-16 | -0.25921 | 0.416 | 0.621 | 5.04E-12 |
| HMGN1 | 1.04E-24 | -0.25943 | 0.178 | 0.369 | 3.39E-20 |
| ZBTB24 | 1.57E-19 | -0.2596 | 0.03 | 0.107 | 5.13E-15 |
| ARID5A | 5.53E-22 | -0.26048 | 0.151 | 0.312 | 1.81E-17 |
| ABRACL | 4.56E-22 | -0.26115 | 0.119 | 0.265 | 1.49E-17 |
| TRAPPC1 | 3.08E-23 | -0.26133 | 0.057 | 0.167 | 1.01E-18 |
| RPL21 | 7.28E-21 | -0.26149 | 0.863 | 0.954 | 2.38E-16 |
| GIMAP7 | 9.18E-28 | -0.26247 | 0.158 | 0.348 | 3.01E-23 |
| ATP5G2 | 7.22E-28 | -0.26249 | 0.262 | 0.508 | 2.37E-23 |
| TMBIM6 | 2.95E-28 | -0.26293 | 0.228 | 0.46 | 9.65E-24 |
| LAMTOR4 | 4.77E-30 | -0.26369 | 0.121 | 0.3 | 1.56E-25 |
| CDKN1A | 7.86E-22 | -0.26407 | 0.111 | 0.249 | 2.57E-17 |
| EEF2 | 7.16E-25 | -0.26466 | 0.321 | 0.584 | 2.35E-20 |
| PPA1 | 7.44E-22 | -0.26517 | 0.085 | 0.208 | 2.44E-17 |
| TOMM20 | 7.48E-27 | -0.2653 | 0.198 | 0.41 | 2.45E-22 |
| FAM107B | 9.17E-18 | -0.26639 | 0.415 | 0.636 | 3.00E-13 |
| PPDPF | 6.51E-26 | -0.26727 | 0.105 | 0.256 | 2.13E-21 |
| SLAMF1 | 5.30E-19 | -0.2673 | 0.061 | 0.159 | 1.74E-14 |
| ATP6V1F | 6.91E-25 | -0.26837 | 0.067 | 0.189 | 2.26E-20 |
| TMEM219 | 2.89E-26 | -0.26839 | 0.058 | 0.179 | 9.46E-22 |
| CLIC1 | 1.05E-27 | -0.26883 | 0.211 | 0.429 | 3.45E-23 |
| CMTM8 | 1.24E-16 | -0.27032 | 0.054 | 0.138 | 4.05E-12 |
| STAM | 2.59E-22 | -0.27097 | 0.07 | 0.185 | 8.48E-18 |
| RPS23 | 5.93E-26 | -0.27148 | 0.858 | 0.949 | 1.94E-21 |
| SRP9 | 9.09E-23 | -0.27225 | 0.061 | 0.171 | 2.97E-18 |
| BTF3 | 4.18E-23 | -0.27333 | 0.403 | 0.666 | 1.37E-18 |
| CDKN1B | 6.06E-25 | -0.27377 | 0.178 | 0.365 | 1.98E-20 |
| IL4I1 | 3.52E-12 | -0.27403 | 0.05 | 0.116 | 1.15E-07 |
| RAC2 | 1.19E-26 | -0.27489 | 0.134 | 0.305 | 3.91E-22 |
| PPP1CB | 1.42E-22 | -0.27516 | 0.31 | 0.549 | 4.65E-18 |
| PAG1 | 3.47E-23 | -0.27572 | 0.241 | 0.447 | 1.14E-18 |
| NDFIP1 | 2.41E-27 | -0.27587 | 0.231 | 0.457 | 7.88E-23 |
| P2RY10 | 1.14E-21 | -0.27598 | 0.089 | 0.213 | 3.74E-17 |
| BRK1 | 2.48E-27 | -0.27626 | 0.09 | 0.236 | 8.12E-23 |
| BAZ1A | 1.65E-20 | -0.27982 | 0.271 | 0.475 | 5.42E-16 |
| RPL37 | 9.57E-24 | -0.28 | 0.807 | 0.942 | 3.13E-19 |
| HIF1A | 1.65E-30 | -0.2805 | 0.118 | 0.293 | 5.41E-26 |
| PRELID1 | 2.52E-28 | -0.28055 | 0.113 | 0.278 | 8.25E-24 |
| KRAS | 5.96E-25 | -0.28062 | 0.135 | 0.3 | 1.95E-20 |
| NAB1 | 1.09E-21 | -0.28068 | 0.04 | 0.13 | 3.56E-17 |
| UBE2D2 | 3.20E-29 | -0.2834 | 0.249 | 0.497 | 1.05E-24 |
| RPL8 | 6.47E-24 | -0.28401 | 0.758 | 0.917 | 2.12E-19 |
| RPL30 | 3.73E-30 | -0.28418 | 0.894 | 0.961 | 1.22E-25 |
| CYTH1 | 1.04E-23 | -0.28439 | 0.368 | 0.63 | 3.40E-19 |
| RHOG | 2.65E-25 | -0.28479 | 0.082 | 0.217 | 8.66E-21 |
| ATRAID | 2.45E-23 | -0.28488 | 0.043 | 0.142 | 8.03E-19 |
| RPL36 | 1.48E-27 | -0.28539 | 0.768 | 0.929 | 4.84E-23 |
| INPP4B | 1.06E-22 | -0.28597 | 0.28 | 0.493 | 3.49E-18 |
| EIF3G | 9.61E-27 | -0.28671 | 0.151 | 0.331 | 3.15E-22 |
| CAPG | 3.00E-22 | -0.2885 | 0.042 | 0.136 | 9.82E-18 |
| ADAM12 | 1.00E-15 | -0.28883 | 0.049 | 0.126 | 3.28E-11 |
| VAMP8 | 4.46E-26 | -0.28914 | 0.11 | 0.265 | 1.46E-21 |
| RPS29 | 6.22E-32 | -0.29315 | 0.898 | 0.971 | 2.04E-27 |
| CD3D | 2.23E-25 | -0.29348 | 0.458 | 0.742 | 7.30E-21 |
| UBA52 | 2.38E-24 | -0.29389 | 0.724 | 0.904 | 7.78E-20 |
| MGAT4A | 1.62E-27 | -0.29569 | 0.2 | 0.407 | 5.31E-23 |
| SPTBN1 | 1.05E-26 | -0.29702 | 0.092 | 0.236 | 3.45E-22 |
| RPLP1 | 4.36E-30 | -0.29753 | 0.945 | 0.984 | 1.43E-25 |
| GPX4 | 3.40E-29 | -0.29853 | 0.162 | 0.36 | 1.11E-24 |
| TMEM14B | 1.89E-26 | -0.29896 | 0.077 | 0.211 | 6.18E-22 |
| TSTD1 | 1.32E-25 | -0.3004 | 0.097 | 0.243 | 4.33E-21 |
| AES | 4.25E-31 | -0.30252 | 0.193 | 0.419 | 1.39E-26 |
| DRAP1 | 1.13E-27 | -0.30254 | 0.127 | 0.298 | 3.71E-23 |
| LRIG1 | 1.11E-22 | -0.30327 | 0.097 | 0.229 | 3.62E-18 |
| SEPW1 | 1.62E-27 | -0.3061 | 0.095 | 0.244 | 5.29E-23 |
| USP3 | 1.57E-28 | -0.30624 | 0.288 | 0.534 | 5.13E-24 |
| RPL10 | 1.63E-33 | -0.30774 | 0.934 | 0.976 | 5.34E-29 |
| RPS8 | 6.28E-29 | -0.30796 | 0.882 | 0.96 | 2.05E-24 |
| RPL37A | 1.92E-28 | -0.30823 | 0.75 | 0.916 | 6.28E-24 |
| RPL32 | 2.07E-32 | -0.30828 | 0.891 | 0.963 | 6.79E-28 |
| RPS9 | 4.82E-27 | -0.30897 | 0.788 | 0.931 | 1.58E-22 |
| YPEL3 | 1.11E-26 | -0.31007 | 0.096 | 0.242 | 3.62E-22 |
| MTMR6 | 1.86E-25 | -0.3104 | 0.06 | 0.178 | 6.08E-21 |
| S100A11 | 6.71E-29 | -0.31041 | 0.389 | 0.658 | 2.20E-24 |
| RPL9 | 7.63E-30 | -0.31046 | 0.82 | 0.946 | 2.50E-25 |
| SNX9 | 1.62E-24 | -0.3111 | 0.256 | 0.468 | 5.32E-20 |
| HNRNPA1 | 2.86E-26 | -0.31113 | 0.428 | 0.696 | 9.35E-22 |
| ARL6IP4 | 3.90E-31 | -0.31165 | 0.147 | 0.342 | 1.28E-26 |
| RPL7A | 4.75E-29 | -0.31216 | 0.813 | 0.937 | 1.55E-24 |
| NDUFS5 | 1.04E-27 | -0.31251 | 0.297 | 0.547 | 3.40E-23 |
| ITPKB | 3.65E-22 | -0.31266 | 0.144 | 0.298 | 1.19E-17 |
| MYL6 | 7.56E-25 | -0.31368 | 0.456 | 0.725 | 2.47E-20 |
| SLCO3A1 | 2.24E-27 | -0.31533 | 0.152 | 0.334 | 7.33E-23 |
| HINT1 | 6.40E-26 | -0.31553 | 0.368 | 0.636 | 2.09E-21 |
| EVI2A | 5.76E-28 | -0.31572 | 0.066 | 0.196 | 1.88E-23 |
| RPL18 | 1.63E-32 | -0.3161 | 0.839 | 0.95 | 5.33E-28 |
| METTL8 | 1.60E-24 | -0.31623 | 0.021 | 0.103 | 5.24E-20 |
| LAT | 1.38E-29 | -0.31817 | 0.067 | 0.203 | 4.53E-25 |
| CD5 | 4.51E-25 | -0.31979 | 0.051 | 0.161 | 1.48E-20 |
| CD48 | 1.15E-28 | -0.31983 | 0.31 | 0.565 | 3.76E-24 |
| MAP3K1 | 1.33E-23 | -0.32048 | 0.086 | 0.213 | 4.34E-19 |
| TLK1 | 4.05E-30 | -0.32068 | 0.212 | 0.432 | 1.33E-25 |
| ARPC3 | 6.87E-31 | -0.32151 | 0.277 | 0.54 | 2.25E-26 |
| PPP1R15A | 3.83E-25 | -0.32168 | 0.134 | 0.296 | 1.25E-20 |
| TMEM243 | 2.29E-30 | -0.32232 | 0.091 | 0.246 | 7.50E-26 |
| TSPO | 2.91E-30 | -0.32237 | 0.138 | 0.325 | 9.54E-26 |
| IL2RG | 7.84E-32 | -0.3228 | 0.179 | 0.396 | 2.57E-27 |
| EIF3K | 2.67E-28 | -0.32311 | 0.267 | 0.519 | 8.73E-24 |
| RPL22 | 4.76E-25 | -0.32331 | 0.62 | 0.851 | 1.56E-20 |
| RUNX1 | 1.88E-28 | -0.32357 | 0.287 | 0.528 | 6.16E-24 |
| TNIK | 5.52E-21 | -0.32452 | 0.351 | 0.581 | 1.81E-16 |
| FRMD4A | 1.26E-20 | -0.3255 | 0.049 | 0.143 | 4.14E-16 |
| TMEM173 | 2.01E-28 | -0.32577 | 0.058 | 0.184 | 6.60E-24 |
| EIF3E | 6.10E-32 | -0.32756 | 0.252 | 0.504 | 2.00E-27 |
| RPL13 | 1.36E-41 | -0.32919 | 0.95 | 0.987 | 4.44E-37 |
| CRTC3 | 1.01E-27 | -0.33054 | 0.077 | 0.215 | 3.32E-23 |
| ISCU | 1.95E-30 | -0.33185 | 0.1 | 0.263 | 6.37E-26 |
| AQP3 | 8.78E-29 | -0.33198 | 0.075 | 0.215 | 2.88E-24 |
| HIVEP1 | 1.41E-26 | -0.3374 | 0.134 | 0.301 | 4.62E-22 |
| PFKFB3 | 1.12E-28 | -0.33749 | 0.131 | 0.306 | 3.66E-24 |
| IRF2BP2 | 1.88E-27 | -0.33759 | 0.139 | 0.313 | 6.16E-23 |
| CIB1 | 1.13E-30 | -0.33802 | 0.207 | 0.431 | 3.68E-26 |
| RPS16 | 1.94E-35 | -0.33834 | 0.839 | 0.951 | 6.36E-31 |
| RPS2 | 2.93E-35 | -0.33838 | 0.889 | 0.964 | 9.59E-31 |
| KDSR | 2.81E-27 | -0.33898 | 0.064 | 0.191 | 9.21E-23 |
| FTH1 | 4.05E-29 | -0.33923 | 0.803 | 0.935 | 1.32E-24 |
| EEF1D | 4.91E-28 | -0.34063 | 0.549 | 0.807 | 1.61E-23 |
| NFATC1 | 1.02E-20 | -0.34109 | 0.044 | 0.135 | 3.35E-16 |
| TNFSF13B | 3.81E-25 | -0.3415 | 0.038 | 0.136 | 1.25E-20 |
| PRDX2 | 2.61E-34 | -0.34266 | 0.094 | 0.265 | 8.53E-30 |
| ZNF331 | 7.84E-24 | -0.34277 | 0.386 | 0.609 | 2.57E-19 |
| RPL27 | 7.17E-31 | -0.34287 | 0.666 | 0.889 | 2.35E-26 |
| RPLP2 | 1.61E-37 | -0.34298 | 0.873 | 0.965 | 5.27E-33 |
| ITM2A | 2.20E-30 | -0.34364 | 0.169 | 0.369 | 7.19E-26 |
| CYTIP | 1.16E-30 | -0.34382 | 0.276 | 0.533 | 3.79E-26 |
| HIST2H2AC | 7.95E-27 | -0.34404 | 0.07 | 0.199 | 2.60E-22 |
| CMSS1 | 9.82E-09 | -0.34441 | 0.229 | 0.339 | 0.000321 |
| ARPC1B | 4.14E-35 | -0.34485 | 0.171 | 0.396 | 1.35E-30 |
| TANK | 2.76E-30 | -0.34663 | 0.231 | 0.459 | 9.03E-26 |
| ESYT2 | 9.08E-33 | -0.34708 | 0.235 | 0.481 | 2.97E-28 |
| ANP32B | 2.29E-35 | -0.35062 | 0.182 | 0.413 | 7.49E-31 |
| GPSM3 | 6.24E-34 | -0.35177 | 0.096 | 0.268 | 2.04E-29 |
| LOH12CR1 | 3.83E-29 | -0.3525 | 0.159 | 0.35 | 1.25E-24 |
| SAMD12 | 7.77E-29 | -0.35613 | 0.091 | 0.24 | 2.54E-24 |
| NACA | 3.89E-31 | -0.35616 | 0.661 | 0.876 | 1.27E-26 |
| APRT | 3.05E-34 | -0.35649 | 0.177 | 0.403 | 9.99E-30 |
| FMN1 | 2.19E-23 | -0.35662 | 0.024 | 0.104 | 7.18E-19 |
| TAF4B | 1.03E-18 | -0.35721 | 0.056 | 0.148 | 3.36E-14 |
| MYL12B | 3.08E-26 | -0.35723 | 0.344 | 0.597 | 1.01E-21 |
| RGS10 | 4.33E-33 | -0.35723 | 0.123 | 0.31 | 1.42E-28 |
| SEPT6 | 1.07E-34 | -0.35728 | 0.275 | 0.54 | 3.49E-30 |
| YWHAB | 4.65E-31 | -0.35729 | 0.242 | 0.483 | 1.52E-26 |
| HIST1H3D | 1.01E-27 | -0.35776 | 0.068 | 0.198 | 3.30E-23 |
| EEF1B2 | 1.95E-28 | -0.35817 | 0.585 | 0.818 | 6.39E-24 |
| RPL15 | 6.59E-33 | -0.35854 | 0.809 | 0.918 | 2.16E-28 |
| SOCS3 | 4.26E-29 | -0.35882 | 0.088 | 0.236 | 1.39E-24 |
| CTLA4 | 2.79E-21 | -0.35926 | 0.039 | 0.127 | 9.13E-17 |
| ABCC1 | 4.18E-34 | -0.35993 | 0.157 | 0.37 | 1.37E-29 |
| RPSA | 1.24E-29 | -0.36171 | 0.656 | 0.875 | 4.07E-25 |
| RPL11 | 3.01E-39 | -0.36216 | 0.876 | 0.965 | 9.87E-35 |
| STAT3 | 3.32E-30 | -0.36227 | 0.284 | 0.532 | 1.09E-25 |
| S1PR1 | 3.91E-29 | -0.36273 | 0.044 | 0.159 | 1.28E-24 |
| EMP3 | 8.05E-33 | -0.36406 | 0.242 | 0.493 | 2.64E-28 |
| GMFG | 2.95E-35 | -0.36479 | 0.258 | 0.523 | 9.66E-31 |
| EID1 | 1.39E-35 | -0.36606 | 0.13 | 0.329 | 4.55E-31 |
| COTL1 | 4.36E-38 | -0.36614 | 0.127 | 0.331 | 1.43E-33 |
| RCAN3 | 3.10E-33 | -0.36777 | 0.204 | 0.434 | 1.01E-28 |
| ICAM2 | 2.69E-25 | -0.36841 | 0.035 | 0.131 | 8.81E-21 |
| SLC25A6 | 4.91E-31 | -0.36875 | 0.256 | 0.502 | 1.61E-26 |
| RPL19 | 6.85E-40 | -0.37031 | 0.88 | 0.961 | 2.24E-35 |
| ARL4C | 1.03E-28 | -0.37087 | 0.425 | 0.693 | 3.38E-24 |
| PHLDB3 | 6.55E-27 | -0.37193 | 0.03 | 0.125 | 2.14E-22 |
| DUSP16 | 2.88E-29 | -0.37241 | 0.256 | 0.49 | 9.41E-25 |
| RPL4 | 6.90E-30 | -0.37259 | 0.45 | 0.72 | 2.26E-25 |
| FAM89B | 2.91E-29 | -0.37351 | 0.059 | 0.187 | 9.51E-25 |
| RPS4X | 4.86E-39 | -0.37383 | 0.88 | 0.959 | 1.59E-34 |
| TMEM123 | 5.67E-33 | -0.37478 | 0.157 | 0.366 | 1.86E-28 |
| CYLD | 1.93E-30 | -0.3752 | 0.145 | 0.336 | 6.33E-26 |
| TAB2 | 3.30E-36 | -0.37546 | 0.173 | 0.396 | 1.08E-31 |
| ITGB1 | 3.36E-32 | -0.37591 | 0.211 | 0.436 | 1.10E-27 |
| LGALS8 | 6.38E-26 | -0.37846 | 0.055 | 0.17 | 2.09E-21 |
| GPCPD1 | 1.54E-24 | -0.37874 | 0.247 | 0.451 | 5.03E-20 |
| BATF | 2.98E-27 | -0.3789 | 0.076 | 0.21 | 9.75E-23 |
| TBL1X | 1.13E-28 | -0.37966 | 0.113 | 0.273 | 3.71E-24 |
| UXS1 | 4.86E-27 | -0.38263 | 0.059 | 0.179 | 1.59E-22 |
| SH3TC1 | 5.39E-32 | -0.38506 | 0.021 | 0.121 | 1.76E-27 |
| AHR | 8.41E-25 | -0.38665 | 0.123 | 0.274 | 2.75E-20 |
| TMEM66 | 7.75E-30 | -0.38793 | 0.579 | 0.832 | 2.54E-25 |
| TMA7 | 8.53E-32 | -0.38833 | 0.329 | 0.603 | 2.79E-27 |
| C11orf31 | 2.45E-30 | -0.39162 | 0.064 | 0.198 | 8.02E-26 |
| CD37 | 3.98E-38 | -0.39249 | 0.189 | 0.431 | 1.30E-33 |
| ARHGAP10 | 1.71E-27 | -0.39322 | 0.072 | 0.204 | 5.61E-23 |
| GLTSCR2 | 1.59E-37 | -0.39453 | 0.289 | 0.567 | 5.22E-33 |
| OCIAD2 | 4.79E-39 | -0.39515 | 0.164 | 0.398 | 1.57E-34 |
| MAST4 | 2.20E-29 | -0.39657 | 0.04 | 0.153 | 7.22E-25 |
| GNB2L1 | 2.56E-33 | -0.39736 | 0.656 | 0.862 | 8.38E-29 |
| RPL5 | 2.05E-35 | -0.40089 | 0.765 | 0.921 | 6.72E-31 |
| ANKRD12 | 2.05E-29 | -0.40299 | 0.534 | 0.792 | 6.71E-25 |
| LEF1 | 6.49E-26 | -0.4066 | 0.03 | 0.124 | 2.12E-21 |
| CNST | 2.94E-33 | -0.4118 | 0.089 | 0.251 | 9.62E-29 |
| RPS18 | 1.34E-48 | -0.41222 | 0.928 | 0.97 | 4.40E-44 |
| FURIN | 4.88E-31 | -0.41245 | 0.033 | 0.142 | 1.60E-26 |
| RPS20 | 1.24E-40 | -0.41418 | 0.716 | 0.909 | 4.05E-36 |
| KIAA1324L | 2.51E-30 | -0.41706 | 0.055 | 0.183 | 8.22E-26 |
| ACTB | 4.04E-39 | -0.41933 | 0.505 | 0.801 | 1.32E-34 |
| TNFRSF18 | 3.06E-30 | -0.41989 | 0.015 | 0.102 | 1.00E-25 |
| PELI1 | 1.08E-30 | -0.42162 | 0.081 | 0.229 | 3.54E-26 |
| RPS3A | 1.50E-44 | -0.42572 | 0.861 | 0.957 | 4.90E-40 |
| NDUFV2 | 2.22E-33 | -0.42617 | 0.151 | 0.353 | 7.27E-29 |
| RPL23 | 3.09E-38 | -0.42854 | 0.578 | 0.834 | 1.01E-33 |
| BTBD11 | 1.61E-23 | -0.43262 | 0.129 | 0.275 | 5.28E-19 |
| FYB | 4.30E-43 | -0.43381 | 0.301 | 0.599 | 1.41E-38 |
| HIST1H1C | 5.38E-35 | -0.43768 | 0.069 | 0.22 | 1.76E-30 |
| IRS2 | 6.65E-34 | -0.44001 | 0.083 | 0.241 | 2.18E-29 |
| TOMM7 | 5.91E-39 | -0.44026 | 0.397 | 0.695 | 1.94E-34 |
| IFITM2 | 5.09E-35 | -0.44115 | 0.257 | 0.507 | 1.67E-30 |
| MAF | 5.68E-28 | -0.44152 | 0.222 | 0.427 | 1.86E-23 |
| ZNRF1 | 1.09E-38 | -0.44325 | 0.079 | 0.25 | 3.58E-34 |
| MYL12A | 5.14E-38 | -0.44338 | 0.304 | 0.586 | 1.68E-33 |
| CD52 | 9.42E-39 | -0.44416 | 0.483 | 0.774 | 3.08E-34 |
| RORA | 1.59E-36 | -0.44461 | 0.471 | 0.754 | 5.21E-32 |
| NDNL2 | 2.04E-35 | -0.4452 | 0.09 | 0.258 | 6.67E-31 |
| RP11-219B17.1 | 1.39E-34 | -0.4453 | 0.194 | 0.417 | 4.56E-30 |
| IL32 | 3.51E-35 | -0.44628 | 0.396 | 0.681 | 1.15E-30 |
| RNASET2 | 4.72E-44 | -0.44691 | 0.154 | 0.396 | 1.55E-39 |
| PRKCA | 1.04E-32 | -0.4498 | 0.212 | 0.434 | 3.41E-28 |
| PTPN13 | 1.26E-33 | -0.45064 | 0.063 | 0.206 | 4.13E-29 |
| EEF1A1 | 2.08E-59 | -0.45196 | 0.934 | 0.986 | 6.80E-55 |
| GNAQ | 3.76E-39 | -0.45249 | 0.078 | 0.25 | 1.23E-34 |
| TRAF1 | 2.65E-37 | -0.45478 | 0.071 | 0.232 | 8.67E-33 |
| MFHAS1 | 4.38E-35 | -0.45544 | 0.047 | 0.181 | 1.43E-30 |
| S100A4 | 9.59E-34 | -0.45941 | 0.514 | 0.779 | 3.14E-29 |
| SERINC5 | 3.12E-41 | -0.45974 | 0.183 | 0.423 | 1.02E-36 |
| RPS6 | 6.21E-50 | -0.46107 | 0.846 | 0.956 | 2.03E-45 |
| EML4 | 5.48E-42 | -0.46182 | 0.55 | 0.832 | 1.79E-37 |
| SPOCK2 | 1.43E-46 | -0.46257 | 0.331 | 0.655 | 4.70E-42 |
| AP3M2 | 1.20E-39 | -0.46385 | 0.102 | 0.291 | 3.94E-35 |
| RPL29 | 1.03E-47 | -0.46621 | 0.74 | 0.924 | 3.37E-43 |
| MZT2A | 1.06E-43 | -0.46896 | 0.143 | 0.374 | 3.46E-39 |
| RPL7 | 3.58E-39 | -0.46916 | 0.616 | 0.857 | 1.17E-34 |
| JAK3 | 1.64E-38 | -0.46927 | 0.049 | 0.193 | 5.38E-34 |
| PLCL1 | 5.11E-30 | -0.46948 | 0.153 | 0.336 | 1.67E-25 |
| CD4 | 2.93E-36 | -0.47119 | 0.061 | 0.21 | 9.58E-32 |
| MAP3K4 | 1.06E-34 | -0.47505 | 0.111 | 0.289 | 3.47E-30 |
| GIMAP4 | 1.14E-39 | -0.47669 | 0.067 | 0.232 | 3.73E-35 |
| CD2 | 1.99E-41 | -0.4772 | 0.42 | 0.727 | 6.50E-37 |
| KLF2 | 8.01E-34 | -0.47954 | 0.332 | 0.577 | 2.62E-29 |
| MBP | 5.30E-42 | -0.48009 | 0.249 | 0.526 | 1.74E-37 |
| AGPAT5 | 1.56E-34 | -0.48092 | 0.048 | 0.179 | 5.12E-30 |
| RPL31 | 2.79E-44 | -0.48433 | 0.565 | 0.849 | 9.14E-40 |
| CLEC2D | 3.17E-50 | -0.48895 | 0.198 | 0.483 | 1.04E-45 |
| TNFAIP8 | 1.06E-45 | -0.49093 | 0.289 | 0.587 | 3.48E-41 |
| SLC2A3 | 8.24E-45 | -0.49111 | 0.276 | 0.562 | 2.70E-40 |
| TNFRSF25 | 5.46E-45 | -0.50139 | 0.077 | 0.264 | 1.79E-40 |
| TBC1D4 | 1.62E-40 | -0.50199 | 0.05 | 0.2 | 5.30E-36 |
| CORO1B | 1.40E-41 | -0.50278 | 0.053 | 0.209 | 4.57E-37 |
| SH3BGRL3 | 2.36E-47 | -0.50583 | 0.448 | 0.771 | 7.73E-43 |
| PFN1 | 6.49E-46 | -0.51131 | 0.398 | 0.714 | 2.13E-41 |
| LIMS1 | 1.68E-43 | -0.51213 | 0.143 | 0.37 | 5.50E-39 |
| DNPH1 | 2.20E-45 | -0.51233 | 0.029 | 0.169 | 7.21E-41 |
| CCR6 | 5.03E-43 | -0.51665 | 0.165 | 0.402 | 1.65E-38 |
| RPL3 | 2.66E-58 | -0.51719 | 0.856 | 0.958 | 8.72E-54 |
| JAZF1 | 2.85E-35 | -0.52694 | 0.201 | 0.422 | 9.34E-31 |
| RHOH | 5.51E-51 | -0.52826 | 0.286 | 0.602 | 1.81E-46 |
| RPS11 | 9.50E-59 | -0.53088 | 0.738 | 0.918 | 3.11E-54 |
| TIAM1 | 5.27E-33 | -0.53735 | 0.023 | 0.127 | 1.72E-28 |
| TPT1 | 1.92E-78 | -0.53761 | 0.927 | 0.981 | 6.30E-74 |
| LDHB | 5.19E-48 | -0.5412 | 0.168 | 0.423 | 1.70E-43 |
| HIST1H1E | 2.18E-44 | -0.5427 | 0.068 | 0.244 | 7.14E-40 |
| CD28 | 6.54E-50 | -0.54466 | 0.086 | 0.293 | 2.14E-45 |
| RPL13A | 4.31E-79 | -0.54696 | 0.883 | 0.978 | 1.41E-74 |
| ARID5B | 5.27E-59 | -0.54886 | 0.259 | 0.583 | 1.73E-54 |
| SELL | 5.70E-39 | -0.55242 | 0.029 | 0.153 | 1.87E-34 |
| BIRC3 | 1.14E-51 | -0.5629 | 0.241 | 0.538 | 3.72E-47 |
| FXYD5 | 2.15E-55 | -0.56807 | 0.226 | 0.533 | 7.03E-51 |
| PASK | 1.44E-48 | -0.56909 | 0.02 | 0.156 | 4.70E-44 |
| ACTG1 | 7.17E-53 | -0.57737 | 0.344 | 0.68 | 2.35E-48 |
| TPM4 | 1.07E-50 | -0.58942 | 0.146 | 0.396 | 3.50E-46 |
| RP4-678D15.1 | 5.71E-45 | -0.61303 | 0.006 | 0.111 | 1.87E-40 |
| CSGALNACT1 | 2.09E-42 | -0.62271 | 0.037 | 0.177 | 6.86E-38 |
| HIST1H1D | 9.36E-49 | -0.63499 | 0.1 | 0.307 | 3.06E-44 |
| GPRIN3 | 4.80E-59 | -0.63825 | 0.209 | 0.509 | 1.57E-54 |
| ADAM19 | 2.52E-43 | -0.63839 | 0.104 | 0.3 | 8.25E-39 |
| MAL | 1.79E-57 | -0.63857 | 0.016 | 0.165 | 5.86E-53 |
| CTSL | 7.46E-48 | -0.63969 | 0.02 | 0.153 | 2.44E-43 |
| RP11-712B9.2 | 5.59E-33 | -0.64496 | 0.041 | 0.162 | 1.83E-28 |
| ETV6 | 7.13E-58 | -0.66379 | 0.156 | 0.427 | 2.33E-53 |
| JUNB | 2.08E-54 | -0.67391 | 0.495 | 0.768 | 6.82E-50 |
| TMEM156 | 1.33E-55 | -0.67965 | 0.048 | 0.234 | 4.34E-51 |
| TNFRSF4 | 1.19E-58 | -0.68185 | 0.02 | 0.177 | 3.89E-54 |
| ZC3H12D | 5.78E-60 | -0.68786 | 0.052 | 0.251 | 1.89E-55 |
| CCR4 | 1.32E-59 | -0.69827 | 0.022 | 0.184 | 4.32E-55 |
| LDLRAD4 | 1.49E-62 | -0.71214 | 0.396 | 0.726 | 4.86E-58 |
| PGAP1 | 7.87E-34 | -0.71241 | 0.093 | 0.251 | 2.58E-29 |
| KLHL5 | 1.34E-50 | -0.72837 | 0.076 | 0.27 | 4.38E-46 |
| TMSB4X | 5.35E-128 | -0.78062 | 0.93 | 0.983 | 1.75E-123 |
| SESN3 | 3.77E-44 | -0.78904 | 0.075 | 0.25 | 1.23E-39 |
| PKIA | 1.27E-68 | -0.83319 | 0.026 | 0.213 | 4.16E-64 |
| PCBP3 | 3.31E-48 | -0.84734 | 0.022 | 0.158 | 1.08E-43 |
| CCR7 | 5.19E-86 | -0.86714 | 0.134 | 0.452 | 1.70E-81 |
| ANK3 | 2.01E-69 | -0.8686 | 0.193 | 0.5 | 6.59E-65 |
| GPR183 | 3.28E-87 | -0.91491 | 0.327 | 0.692 | 1.07E-82 |
| ICOS | 2.05E-102 | -1.05807 | 0.179 | 0.555 | 6.72E-98 |
| RPS26 | 6.26E-141 | -1.18672 | 0.443 | 0.818 | 2.05E-136 |
| LTB | 8.95E-153 | -1.52955 | 0.196 | 0.646 | 2.93E-148 |
| TSHZ2 | 7.43E-131 | -1.79594 | 0.043 | 0.366 | 2.43E-126 |
| FAAH2 | 1.09E-124 | -1.82941 | 0.074 | 0.412 | 3.57E-120 |

**Supplementary Table 4: Top significant DE genes comparing expected tissue-resident and blood NK cells in PDAC tumor tissue, as identified by Originator. log2 fold change (avg_log2FC) is averaged expression in expected tissue-resident compared to blood NK cells.**

|  | **p_val** | **avg_log2FC** | **pct.1** | **pct.2** | **p_val_adj** |
| --- | --- | --- | --- | --- | --- |
| MLLT3 | 1.58E-08 | 0.919308 | 0.58 | 0.378 | 0.000517 |
| GZMK | 4.42E-17 | 0.856396 | 0.812 | 0.459 | 1.45E-12 |
| CCDC64 | 7.68E-09 | 0.825476 | 0.567 | 0.369 | 0.000251 |
| RGCC | 1.47E-10 | 0.792257 | 0.778 | 0.617 | 4.82E-06 |
| ZNF331 | 9.65E-08 | 0.632912 | 0.776 | 0.635 | 0.003159 |
| CD44 | 7.81E-08 | 0.528318 | 0.835 | 0.739 | 0.002555 |
| TMSB4X | 4.69E-08 | -0.35113 | 0.998 | 1 | 0.001535 |
| SH3BGRL3 | 1.85E-07 | -0.4571 | 0.724 | 0.865 | 0.006045 |
| PFN1 | 1.68E-07 | -0.47928 | 0.678 | 0.842 | 0.005509 |
| TMSB10 | 3.19E-08 | -0.48381 | 0.906 | 0.955 | 0.001044 |
| ARPC2 | 3.52E-10 | -0.60215 | 0.561 | 0.797 | 1.15E-05 |
| HCST | 1.66E-08 | -0.60324 | 0.516 | 0.725 | 0.000545 |
| MIR4435-1HG | 2.60E-08 | -0.60849 | 0.108 | 0.275 | 0.000853 |
| SYNGR1 | 6.01E-08 | -0.62617 | 0.02 | 0.113 | 0.001966 |
| CHST12 | 1.17E-08 | -0.645 | 0.082 | 0.239 | 0.000382 |
| MYL12A | 1.87E-10 | -0.71587 | 0.467 | 0.689 | 6.11E-06 |
| ACTG1 | 8.24E-09 | -0.73712 | 0.516 | 0.716 | 0.00027 |
| ITGB1 | 4.16E-08 | -0.73913 | 0.247 | 0.441 | 0.001361 |
| LYN | 6.91E-09 | -0.77626 | 0.027 | 0.14 | 0.000226 |
| CD63 | 1.91E-10 | -0.77783 | 0.194 | 0.419 | 6.24E-06 |
| KLRF1 | 4.79E-09 | -0.8157 | 0.027 | 0.14 | 0.000157 |
| CTB-91J4.1 | 7.64E-09 | -0.83219 | 0.02 | 0.122 | 0.00025 |
| EFHD2 | 1.38E-10 | -0.84341 | 0.2 | 0.432 | 4.52E-06 |
| ACTB | 4.75E-15 | -0.87106 | 0.69 | 0.856 | 1.55E-10 |
| AC092580.4 | 7.47E-11 | -0.87745 | 0.112 | 0.311 | 2.44E-06 |
| NCALD | 3.63E-10 | -0.92863 | 0.1 | 0.279 | 1.19E-05 |
| AGPAT4 | 1.02E-14 | -1.10832 | 0.112 | 0.351 | 3.33E-10 |
| HOPX | 1.16E-17 | -1.14243 | 0.063 | 0.297 | 3.81E-13 |
| KLRD1 | 1.25E-18 | -1.17805 | 0.106 | 0.378 | 4.08E-14 |
| LGALS1 | 3.51E-12 | -1.20687 | 0.216 | 0.455 | 1.15E-07 |
| KLRB1 | 1.46E-11 | -1.23468 | 0.076 | 0.257 | 4.77E-07 |
| LINGO2 | 4.02E-08 | -1.24521 | 0.016 | 0.104 | 0.001316 |
| TYROBP | 2.03E-16 | -1.24536 | 0.051 | 0.257 | 6.65E-12 |
| S100A4 | 1.10E-18 | -1.31244 | 0.365 | 0.676 | 3.61E-14 |
| FCGR3A | 2.51E-20 | -1.37518 | 0.02 | 0.221 | 8.20E-16 |
| NKG7 | 2.64E-40 | -1.46565 | 0.569 | 0.923 | 8.63E-36 |
| GZMH | 5.90E-35 | -1.55084 | 0.224 | 0.689 | 1.93E-30 |
| FGFBP2 | 7.79E-20 | -1.58029 | 0.041 | 0.27 | 2.55E-15 |
| PRF1 | 8.19E-24 | -1.62711 | 0.09 | 0.396 | 2.68E-19 |
| GZMB | 1.77E-41 | -2.31745 | 0.063 | 0.486 | 5.79E-37 |
| GNLY | 1.07E-41 | -2.91977 | 0.065 | 0.486 | 3.50E-37 |

**Supplementary Table 5: Top significant DE genes comparing expected tissue-resident and blood macrophage cells in PDAC tumor tissue, as identified by Originator. log2 fold change (avg_log2FC) is averaged expression in expected tissue-resident compared to blood macrophage cells.**

|  | **p_val** | **avg_log2FC** | **pct.1** | **pct.2** | **p_val_adj** |
| --- | --- | --- | --- | --- | --- |
| EREG | 5.77E-12 | 1.587125 | 0.455 | 0.302 | 1.89E-07 |
| RND3 | 2.22E-19 | 1.572774 | 0.521 | 0.314 | 7.27E-15 |
| TNF | 1.48E-34 | 1.40944 | 0.658 | 0.314 | 4.84E-30 |
| CXCL3 | 1.52E-30 | 1.337026 | 0.819 | 0.644 | 4.99E-26 |
| LSAMP | 1.36E-20 | 1.330233 | 0.423 | 0.173 | 4.45E-16 |
| INHBA | 3.15E-13 | 1.308891 | 0.597 | 0.461 | 1.03E-08 |
| CCL20 | 1.11E-15 | 1.301512 | 0.496 | 0.292 | 3.62E-11 |
| FABP4 | 3.33E-21 | 1.269431 | 0.377 | 0.132 | 1.09E-16 |
| SGMS2 | 7.79E-22 | 1.191039 | 0.631 | 0.434 | 2.55E-17 |
| IL1B | 9.05E-19 | 1.185804 | 0.787 | 0.619 | 2.96E-14 |
| TEX14 | 3.65E-16 | 1.132171 | 0.736 | 0.551 | 1.19E-11 |
| ANXA1 | 1.47E-24 | 1.085911 | 0.941 | 0.901 | 4.80E-20 |
| ANKRD28 | 5.87E-32 | 1.056255 | 0.829 | 0.64 | 1.92E-27 |
| SVIL | 7.72E-19 | 1.00178 | 0.562 | 0.347 | 2.53E-14 |
| JUN | 3.14E-28 | 0.940978 | 0.966 | 0.892 | 1.03E-23 |
| DNAJB1 | 9.50E-27 | 0.934928 | 0.858 | 0.65 | 3.11E-22 |
| IER2 | 5.51E-25 | 0.932401 | 0.939 | 0.828 | 1.80E-20 |
| EGR1 | 1.30E-28 | 0.923217 | 0.895 | 0.67 | 4.27E-24 |
| B3GNT5 | 1.01E-26 | 0.923103 | 0.807 | 0.625 | 3.30E-22 |
| LRRC23 | 5.99E-11 | 0.915788 | 0.494 | 0.324 | 1.96E-06 |
| CITED2 | 4.70E-27 | 0.911896 | 0.778 | 0.585 | 1.54E-22 |
| KCNQ1OT1 | 2.90E-08 | 0.908648 | 0.345 | 0.209 | 0.00095 |
| PPP1R15A | 8.98E-35 | 0.905377 | 0.936 | 0.797 | 2.94E-30 |
| DENND5A | 9.17E-23 | 0.905237 | 0.848 | 0.788 | 3.00E-18 |
| IL1A | 3.74E-16 | 0.895255 | 0.565 | 0.366 | 1.22E-11 |
| BTG2 | 3.11E-29 | 0.893733 | 0.907 | 0.776 | 1.02E-24 |
| ADAM17 | 1.11E-35 | 0.885103 | 0.902 | 0.797 | 3.62E-31 |
| NFKBIZ | 1.34E-31 | 0.863318 | 0.941 | 0.756 | 4.40E-27 |
| OASL | 2.52E-24 | 0.861963 | 0.504 | 0.237 | 8.25E-20 |
| KLF4 | 5.26E-30 | 0.860441 | 0.936 | 0.797 | 1.72E-25 |
| CXCL2 | 1.37E-15 | 0.856468 | 0.807 | 0.655 | 4.48E-11 |
| DUSP2 | 3.87E-21 | 0.854534 | 0.802 | 0.599 | 1.27E-16 |
| FOS | 7.55E-24 | 0.838205 | 0.966 | 0.901 | 2.47E-19 |
| EIF4E | 3.43E-11 | 0.826687 | 0.67 | 0.561 | 1.12E-06 |
| HSPA1B | 5.91E-17 | 0.816766 | 0.746 | 0.554 | 1.93E-12 |
| CCL3 | 1.11E-09 | 0.810991 | 0.885 | 0.836 | 3.62E-05 |
| HSPA1A | 1.58E-13 | 0.804674 | 0.897 | 0.788 | 5.17E-09 |
| KB-1507C5.4 | 8.71E-10 | 0.794849 | 0.565 | 0.415 | 2.85E-05 |
| RAB7A | 3.56E-25 | 0.791889 | 0.973 | 0.921 | 1.17E-20 |
| UAP1 | 2.85E-20 | 0.791087 | 0.575 | 0.345 | 9.33E-16 |
| PMAIP1 | 3.82E-09 | 0.769728 | 0.562 | 0.424 | 0.000125 |
| SIK2 | 1.55E-07 | 0.768383 | 0.65 | 0.573 | 0.005088 |
| ATP2B1 | 3.08E-16 | 0.76712 | 0.922 | 0.896 | 1.01E-11 |
| PHLDA1 | 2.51E-23 | 0.756932 | 0.758 | 0.523 | 8.20E-19 |
| ADIPOR2 | 1.73E-10 | 0.755075 | 0.641 | 0.575 | 5.66E-06 |
| B4GALT1 | 9.45E-24 | 0.752053 | 0.902 | 0.818 | 3.10E-19 |
| DNAJA1 | 7.43E-25 | 0.751925 | 0.958 | 0.889 | 2.43E-20 |
| RGCC | 1.04E-07 | 0.743094 | 0.719 | 0.64 | 0.003401 |
| SEMA4A | 1.26E-09 | 0.741826 | 0.709 | 0.596 | 4.13E-05 |
| KLHL29 | 2.08E-18 | 0.741634 | 0.384 | 0.156 | 6.82E-14 |
| MMP19 | 4.95E-12 | 0.741109 | 0.689 | 0.541 | 1.62E-07 |
| MCF2L2 | 1.50E-29 | 0.731839 | 0.609 | 0.298 | 4.92E-25 |
| SHROOM3 | 2.05E-12 | 0.7092 | 0.174 | 0.046 | 6.71E-08 |
| RP11-701P16.5 | 3.90E-15 | 0.708258 | 0.484 | 0.281 | 1.28E-10 |
| ATP10A | 9.29E-16 | 0.707077 | 0.394 | 0.178 | 3.04E-11 |
| IL1RN | 1.59E-07 | 0.702058 | 0.575 | 0.465 | 0.00519 |
| FOSB | 4.48E-25 | 0.700165 | 0.985 | 0.853 | 1.47E-20 |
| C19orf59 | 1.71E-20 | 0.697194 | 0.535 | 0.274 | 5.59E-16 |
| RP11-779O18.3 | 1.58E-19 | 0.696941 | 0.733 | 0.479 | 5.18E-15 |
| ZSWIM6 | 8.47E-19 | 0.695827 | 0.985 | 0.941 | 2.77E-14 |
| DUSP1 | 9.54E-24 | 0.692711 | 0.973 | 0.899 | 3.12E-19 |
| DNAJA4 | 9.30E-17 | 0.684884 | 0.491 | 0.274 | 3.04E-12 |
| INTS6 | 5.41E-11 | 0.683984 | 0.672 | 0.529 | 1.77E-06 |
| IL8 | 2.08E-12 | 0.66944 | 0.902 | 0.772 | 6.81E-08 |
| CLIC4 | 1.57E-13 | 0.667582 | 0.885 | 0.779 | 5.14E-09 |
| HDAC9 | 1.17E-07 | 0.663882 | 0.34 | 0.209 | 0.003832 |
| KLF2 | 3.05E-14 | 0.656353 | 0.795 | 0.618 | 9.99E-10 |
| TNFAIP6 | 5.24E-11 | 0.656306 | 0.333 | 0.176 | 1.72E-06 |
| HBEGF | 1.91E-09 | 0.649177 | 0.697 | 0.607 | 6.25E-05 |
| MACC1 | 2.82E-07 | 0.63612 | 0.445 | 0.33 | 0.009235 |
| TCOF1 | 1.83E-11 | 0.624082 | 0.623 | 0.465 | 5.99E-07 |
| LINC00936 | 4.31E-19 | 0.618987 | 0.658 | 0.43 | 1.41E-14 |
| SERTAD1 | 5.26E-18 | 0.618646 | 0.724 | 0.529 | 1.72E-13 |
| CDKN1A | 8.61E-09 | 0.617549 | 0.68 | 0.575 | 0.000282 |
| FBXO11 | 1.43E-15 | 0.615544 | 0.878 | 0.785 | 4.69E-11 |
| RP11-58E21.3 | 6.46E-13 | 0.610365 | 0.32 | 0.147 | 2.12E-08 |
| NR4A1 | 5.00E-17 | 0.605293 | 0.785 | 0.609 | 1.64E-12 |
| UBB | 3.79E-21 | 0.601623 | 0.99 | 0.969 | 1.24E-16 |
| TREM1 | 6.89E-11 | 0.59822 | 0.697 | 0.615 | 2.26E-06 |
| ATF3 | 2.93E-20 | 0.598132 | 0.949 | 0.773 | 9.60E-16 |
| MIR155HG | 7.26E-09 | 0.592086 | 0.663 | 0.51 | 0.000238 |
| HSPH1 | 9.43E-08 | 0.590248 | 0.773 | 0.695 | 0.003088 |
| NFKBIA | 3.63E-17 | 0.589396 | 0.993 | 0.919 | 1.19E-12 |
| PPP1R10 | 2.82E-15 | 0.582846 | 0.724 | 0.544 | 9.23E-11 |
| TNFRSF10B | 2.21E-09 | 0.576509 | 0.579 | 0.439 | 7.22E-05 |
| H3F3B | 1.64E-25 | 0.573426 | 0.995 | 0.979 | 5.38E-21 |
| TUBB4B | 1.06E-11 | 0.572193 | 0.814 | 0.708 | 3.46E-07 |
| ABL2 | 4.67E-13 | 0.571107 | 0.883 | 0.806 | 1.53E-08 |
| EGR2 | 5.35E-10 | 0.569174 | 0.667 | 0.498 | 1.75E-05 |
| TSC22D2 | 4.97E-17 | 0.56705 | 0.897 | 0.742 | 1.63E-12 |
| OTUD1 | 1.22E-13 | 0.566881 | 0.653 | 0.477 | 3.99E-09 |
| KCNE1 | 1.24E-11 | 0.562304 | 0.553 | 0.361 | 4.08E-07 |
| TCF7L2 | 1.02E-14 | 0.56188 | 0.775 | 0.639 | 3.35E-10 |
| STK17B | 5.56E-16 | 0.550407 | 0.638 | 0.437 | 1.82E-11 |
| RP11-24F11.2 | 9.95E-12 | 0.548366 | 0.391 | 0.227 | 3.26E-07 |
| IGFBP2 | 5.48E-08 | 0.543188 | 0.433 | 0.287 | 0.001793 |
| CCDC18 | 2.78E-09 | 0.541281 | 0.455 | 0.299 | 9.10E-05 |
| UBE2R2 | 2.45E-09 | 0.53771 | 0.773 | 0.708 | 8.01E-05 |
| NFE2L3 | 1.81E-09 | 0.536286 | 0.499 | 0.332 | 5.94E-05 |
| RP11-598F7.3 | 8.00E-09 | 0.534108 | 0.423 | 0.29 | 0.000262 |

**Supplementary Table 6: Top significant DE genes comparing expected tissue-resident and blood monocytes in PDAC tumor tissue, as identified by Originator. log2 fold change (avg_log2FC) is averaged expression in expected tissue-resident compared to blood monocytes.**

|  | p_val | avg_log2FC | pct.1 | pct.2 | p_val_adj |
| --- | --- | --- | --- | --- | --- |
| CPA3 | 3.10E-15 | 1.760257 | 0.2 | 0 | 1.02E-10 |
| KRT5 | 1.46E-19 | 1.741882 | 0.4 | 0.003 | 4.78E-15 |
| CLEC4C | 2.42E-12 | 1.737032 | 0.4 | 0.01 | 7.93E-08 |
| PACSIN1 | 1.82E-10 | 1.715609 | 0.4 | 0.013 | 5.95E-06 |
| LILRA5 | 6.18E-09 | 1.563429 | 0.8 | 0.08 | 0.000202 |
| SCT | 1.82E-10 | 1.526098 | 0.4 | 0.013 | 5.95E-06 |
| TPSAB1 | 3.51E-08 | 1.365937 | 0.2 | 0.003 | 0.001149 |
| VASH2 | 3.35E-12 | 1.357507 | 0.4 | 0.01 | 1.10E-07 |
| MAP1A | 5.07E-08 | 1.336734 | 0.4 | 0.019 | 0.001658 |
| CERS4 | 1.49E-09 | 1.221591 | 0.6 | 0.038 | 4.86E-05 |
| LILRA1 | 4.02E-09 | 1.124013 | 0.4 | 0.016 | 0.000131 |
| ASIP | 4.13E-08 | 1.101175 | 0.4 | 0.019 | 0.001352 |
| CTD-2023N9.1 | 4.62E-29 | 1.029141 | 0.4 | 0 | 1.51E-24 |
| BSPRY | 8.53E-20 | 0.847571 | 0.4 | 0.003 | 2.79E-15 |
| TAL1 | 3.10E-15 | 0.844676 | 0.2 | 0 | 1.02E-10 |
| AC097495.2 | 3.10E-15 | 0.844676 | 0.2 | 0 | 1.02E-10 |
| LCN6.1 | 3.10E-15 | 0.844676 | 0.2 | 0 | 1.02E-10 |
| RP11-573G6.10 | 3.10E-15 | 0.844676 | 0.2 | 0 | 1.02E-10 |
| OR10A4 | 3.10E-15 | 0.844676 | 0.2 | 0 | 1.02E-10 |
| EMID1 | 3.10E-15 | 0.844676 | 0.2 | 0 | 1.02E-10 |
| PPARGC1A | 3.51E-08 | 0.841151 | 0.2 | 0.003 | 0.001149 |
| F2RL3 | 3.51E-08 | 0.836686 | 0.2 | 0.003 | 0.001149 |
| AC074117.10 | 4.02E-09 | 0.831836 | 0.4 | 0.016 | 0.000131 |
| SMPD3 | 3.94E-12 | 0.819265 | 0.4 | 0.01 | 1.29E-07 |
| LRRC36 | 3.51E-08 | 0.812328 | 0.2 | 0.003 | 0.001149 |
| PTGES | 5.73E-15 | 0.568231 | 0.4 | 0.006 | 1.87E-10 |
| N4BP3 | 5.07E-08 | 0.555367 | 0.4 | 0.019 | 0.001658 |
| AC062017.1 | 9.27E-08 | 0.537653 | 0.4 | 0.019 | 0.003033 |
| RIMKLA | 3.10E-15 | 0.509674 | 0.2 | 0 | 1.02E-10 |
| RP11-161D15.3 | 3.10E-15 | 0.509674 | 0.2 | 0 | 1.02E-10 |
| RP11-844P9.2 | 3.10E-15 | 0.509674 | 0.2 | 0 | 1.02E-10 |
| GATSL1 | 3.10E-15 | 0.509674 | 0.2 | 0 | 1.02E-10 |
| ATE1-AS1 | 3.10E-15 | 0.509674 | 0.2 | 0 | 1.02E-10 |
| ADAMTSL3 | 3.10E-15 | 0.509674 | 0.2 | 0 | 1.02E-10 |
| C19orf84 | 3.10E-15 | 0.509674 | 0.2 | 0 | 1.02E-10 |
| AC004837.5 | 3.51E-08 | 0.507871 | 0.2 | 0.003 | 0.001149 |
| RP11-783K16.14 | 3.51E-08 | 0.507136 | 0.2 | 0.003 | 0.001149 |
| RP11-530C5.1 | 3.51E-08 | 0.507107 | 0.2 | 0.003 | 0.001149 |
| RP11-473O4.5 | 3.51E-08 | 0.506426 | 0.2 | 0.003 | 0.001149 |
| RP11-275F13.1 | 3.51E-08 | 0.505324 | 0.2 | 0.003 | 0.001149 |
| RP11-134O21.1 | 3.51E-08 | 0.504989 | 0.2 | 0.003 | 0.001149 |
| FAM227A | 3.51E-08 | 0.501919 | 0.2 | 0.003 | 0.001149 |
| FAM69B | 3.51E-08 | 0.500147 | 0.2 | 0.003 | 0.001149 |
| RP11-71G12.1 | 4.30E-08 | 0.494773 | 0.2 | 0.003 | 0.001407 |
| AC011893.3 | 3.10E-15 | 0.464982 | 0.2 | 0 | 1.02E-10 |
| RP11-33O4.1 | 3.10E-15 | 0.464982 | 0.2 | 0 | 1.02E-10 |
| GP5 | 3.10E-15 | 0.464982 | 0.2 | 0 | 1.02E-10 |
| SBSPON | 3.10E-15 | 0.464982 | 0.2 | 0 | 1.02E-10 |
| RP11-388P9.2 | 3.10E-15 | 0.464982 | 0.2 | 0 | 1.02E-10 |
| KCNA5 | 3.10E-15 | 0.464982 | 0.2 | 0 | 1.02E-10 |
| CTD-2292M16.8 | 3.10E-15 | 0.464982 | 0.2 | 0 | 1.02E-10 |
| LTK | 3.10E-15 | 0.464982 | 0.2 | 0 | 1.02E-10 |
| MAG | 3.10E-15 | 0.464982 | 0.2 | 0 | 1.02E-10 |
| AP001439.2 | 3.10E-15 | 0.464982 | 0.2 | 0 | 1.02E-10 |
| RP11-388M20.1 | 3.51E-08 | 0.461734 | 0.2 | 0.003 | 0.001149 |
| ARHGEF16 | 3.51E-08 | 0.461418 | 0.2 | 0.003 | 0.001149 |
| RP5-1031D4.2 | 3.51E-08 | 0.461243 | 0.2 | 0.003 | 0.001149 |
| WNT10A | 3.51E-08 | 0.457512 | 0.2 | 0.003 | 0.001149 |
| PPFIA4 | 3.10E-15 | 0.318361 | 0.2 | 0 | 1.02E-10 |
| RP11-31K23.2 | 3.51E-08 | 0.316219 | 0.2 | 0.003 | 0.001149 |
| RP11-505P4.6 | 3.10E-15 | 0.316085 | 0.2 | 0 | 1.02E-10 |
| AC020629.1 | 3.10E-15 | 0.316085 | 0.2 | 0 | 1.02E-10 |
| PRKD1 | 3.51E-08 | 0.315562 | 0.2 | 0.003 | 0.001149 |
| KCNH3 | 3.51E-08 | 0.314667 | 0.2 | 0.003 | 0.001149 |
| CTD-2267D19.3 | 3.51E-08 | 0.314257 | 0.2 | 0.003 | 0.001149 |
| AC007970.1 | 3.51E-08 | 0.313992 | 0.2 | 0.003 | 0.001149 |
| RP11-290L1.3 | 3.51E-08 | 0.313951 | 0.2 | 0.003 | 0.001149 |
| CLMP | 3.51E-08 | 0.313762 | 0.2 | 0.003 | 0.001149 |
| SHPK | 3.51E-08 | 0.312722 | 0.2 | 0.003 | 0.001149 |
| SEZ6 | 3.51E-08 | 0.311999 | 0.2 | 0.003 | 0.001149 |
| SERPINB10 | 4.30E-08 | 0.306728 | 0.2 | 0.003 | 0.001407 |

**Supplementary Table 7: Top significant DE genes comparing fetal and maternal fibroblast type 1 cells in placenta tissue, as identified by Originator. log2 fold change (avg_log2FC) is averaged expression in fetal compared to maternal fibroblast type 1.**

|  | **p_val** | **avg_log2FC** | **pct.1** | **pct.2** | **p_val_adj** |
| --- | --- | --- | --- | --- | --- |
| EGFL6 | 1.59E-243 | 4.429656 | 0.961 | 0.125 | 3.51E-239 |
| TCF21 | 1.86E-197 | 4.315008 | 0.868 | 0.089 | 4.13E-193 |
| GPC3 | 3.45E-235 | 4.239064 | 0.946 | 0.114 | 7.65E-231 |
| PLA2G2A | 8.73E-156 | 4.072042 | 0.79 | 0.105 | 1.93E-151 |
| CD36 | 5.40E-180 | 3.637041 | 0.831 | 0.093 | 1.20E-175 |
| WNT2 | 1.56E-154 | 3.371118 | 0.753 | 0.039 | 3.46E-150 |
| HSD17B2 | 1.30E-197 | 3.30591 | 0.883 | 0.155 | 2.88E-193 |
| DLK1 | 1.10E-162 | 3.288633 | 0.817 | 0.098 | 2.43E-158 |
| C7 | 1.69E-138 | 3.251991 | 0.748 | 0.095 | 3.75E-134 |
| HAPLN1 | 3.37E-91 | 3.108324 | 0.545 | 0.011 | 7.47E-87 |
| RARRES2 | 2.42E-143 | 3.066998 | 0.757 | 0.091 | 5.37E-139 |
| PITX2 | 7.50E-145 | 3.029137 | 0.724 | 0.03 | 1.66E-140 |
| PHACTR2 | 3.62E-166 | 2.886097 | 0.827 | 0.202 | 8.02E-162 |
| HGF | 3.26E-101 | 2.754256 | 0.606 | 0.05 | 7.24E-97 |
| SEPP1 | 1.24E-175 | 2.721012 | 0.936 | 0.673 | 2.75E-171 |
| MEG3 | 1.40E-212 | 2.720558 | 0.981 | 0.659 | 3.10E-208 |
| SERPINF1 | 1.88E-171 | 2.709533 | 0.929 | 0.641 | 4.18E-167 |
| PMP22 | 2.47E-162 | 2.605238 | 0.864 | 0.352 | 5.47E-158 |
| COL3A1 | 4.15E-149 | 2.567016 | 0.891 | 0.475 | 9.19E-145 |
| CYTL1 | 7.92E-54 | 2.556446 | 0.378 | 0.005 | 1.75E-49 |
| C12orf39 | 3.08E-56 | 2.458759 | 0.396 | 0.009 | 6.83E-52 |
| CNN3 | 7.64E-148 | 2.448055 | 0.825 | 0.227 | 1.69E-143 |
| RPS23 | 1.09E-226 | 2.393955 | 0.997 | 0.827 | 2.42E-222 |
| S100A10 | 1.78E-160 | 2.389769 | 0.891 | 0.373 | 3.95E-156 |
| FGF7 | 7.51E-54 | 2.342571 | 0.488 | 0.152 | 1.67E-49 |
| TAGLN2 | 1.63E-120 | 2.290443 | 0.8 | 0.395 | 3.61E-116 |
| GPX3 | 2.58E-108 | 2.251919 | 0.812 | 0.361 | 5.72E-104 |
| ENPP2 | 2.21E-112 | 2.238685 | 0.77 | 0.33 | 4.90E-108 |
| TFPI2 | 1.04E-32 | 2.16592 | 0.321 | 0.061 | 2.30E-28 |
| PDPN | 1.89E-99 | 2.126445 | 0.791 | 0.573 | 4.20E-95 |
| CADM3 | 2.79E-65 | 2.109031 | 0.438 | 0.011 | 6.18E-61 |
| MATN2 | 1.01E-84 | 2.108842 | 0.632 | 0.216 | 2.25E-80 |
| AKR1B1 | 8.04E-72 | 2.067901 | 0.742 | 0.58 | 1.78E-67 |
| ANGPTL1 | 3.19E-85 | 2.058844 | 0.633 | 0.18 | 7.08E-81 |
| TMEM176A | 3.96E-80 | 2.052475 | 0.646 | 0.223 | 8.78E-76 |
| RPL23A | 4.90E-210 | 2.044697 | 0.997 | 0.814 | 1.09E-205 |
| PLAC1 | 9.13E-62 | 2.017424 | 0.424 | 0.014 | 2.02E-57 |
| SORBS2 | 1.47E-75 | 1.993688 | 0.513 | 0.043 | 3.26E-71 |
| RPL27A | 1.57E-213 | 1.992075 | 0.998 | 0.88 | 3.49E-209 |
| RPS27 | 6.63E-175 | 1.947134 | 0.986 | 0.752 | 1.47E-170 |
| RPL36A | 2.07E-123 | 1.85864 | 0.845 | 0.45 | 4.59E-119 |
| RPL21 | 1.80E-184 | 1.858082 | 0.994 | 0.814 | 4.00E-180 |
| RPL26 | 1.14E-191 | 1.844048 | 0.997 | 0.857 | 2.53E-187 |
| RPL34 | 1.75E-180 | 1.843375 | 0.995 | 0.83 | 3.87E-176 |
| RPS25 | 9.10E-160 | 1.827714 | 0.964 | 0.645 | 2.02E-155 |
| RPS28 | 3.29E-152 | 1.816583 | 0.949 | 0.695 | 7.30E-148 |
| BMP5 | 9.49E-59 | 1.802208 | 0.414 | 0.016 | 2.10E-54 |
| RPS15A | 1.64E-178 | 1.793451 | 0.99 | 0.782 | 3.62E-174 |
| RPL7 | 7.21E-193 | 1.791191 | 0.997 | 0.868 | 1.60E-188 |
| BMP4 | 6.32E-52 | 1.77684 | 0.424 | 0.068 | 1.40E-47 |
| TMEM176B | 1.23E-87 | 1.759691 | 0.776 | 0.386 | 2.72E-83 |
| RPS3A | 8.65E-179 | 1.747539 | 0.991 | 0.825 | 1.92E-174 |
| RPL31 | 7.48E-169 | 1.733673 | 0.993 | 0.789 | 1.66E-164 |
| RPL38 | 9.06E-119 | 1.720346 | 0.863 | 0.516 | 2.01E-114 |
| CD9 | 4.82E-76 | 1.714701 | 0.653 | 0.218 | 1.07E-71 |
| NUPR1 | 6.29E-97 | 1.695084 | 0.85 | 0.673 | 1.39E-92 |
| RPS14 | 9.13E-174 | 1.663032 | 0.998 | 0.907 | 2.02E-169 |
| FRZB | 2.03E-30 | 1.655166 | 0.31 | 0.057 | 4.51E-26 |
| COL15A1 | 1.55E-43 | 1.653874 | 0.46 | 0.164 | 3.44E-39 |
| RPS4Y1 | 4.71E-14 | 1.652484 | 0.314 | 0.193 | 1.05E-09 |
| RPS27A | 1.26E-175 | 1.651167 | 0.996 | 0.902 | 2.79E-171 |
| RPS29 | 2.65E-88 | 1.649661 | 0.754 | 0.373 | 5.88E-84 |
| RPL30 | 1.02E-159 | 1.646104 | 0.99 | 0.791 | 2.26E-155 |
| HNRNPA1 | 1.96E-107 | 1.633338 | 0.838 | 0.625 | 4.34E-103 |
| RPS24 | 5.74E-160 | 1.631131 | 0.986 | 0.805 | 1.27E-155 |
| FIBIN | 6.17E-42 | 1.624765 | 0.333 | 0.02 | 1.37E-37 |
| UQCRB | 1.13E-54 | 1.618989 | 0.642 | 0.38 | 2.49E-50 |
| GNG11 | 1.96E-32 | 1.587127 | 0.532 | 0.339 | 4.34E-28 |
| RPL24 | 2.99E-160 | 1.577585 | 0.981 | 0.82 | 6.63E-156 |
| RPL14 | 1.25E-164 | 1.569504 | 0.991 | 0.877 | 2.77E-160 |
| SEPT7 | 6.35E-75 | 1.569425 | 0.731 | 0.505 | 1.41E-70 |
| SPARC | 1.70E-171 | 1.564863 | 0.99 | 0.92 | 3.77E-167 |
| RPLP2 | 7.57E-162 | 1.545529 | 0.997 | 0.886 | 1.68E-157 |
| EIF3E | 8.54E-76 | 1.543778 | 0.759 | 0.548 | 1.89E-71 |
| NPM1 | 1.01E-120 | 1.539533 | 0.892 | 0.716 | 2.23E-116 |
| BST2 | 6.73E-49 | 1.514151 | 0.584 | 0.27 | 1.49E-44 |
| ELN | 2.76E-38 | 1.502515 | 0.342 | 0.048 | 6.12E-34 |
| RPL15 | 7.38E-176 | 1.49784 | 0.998 | 0.964 | 1.64E-171 |
| RPS8 | 5.67E-154 | 1.496914 | 0.997 | 0.893 | 1.26E-149 |
| PCOLCE2 | 4.58E-45 | 1.496335 | 0.371 | 0.036 | 1.01E-40 |
| PLP2 | 1.86E-55 | 1.493829 | 0.652 | 0.436 | 4.13E-51 |
| PTGDS | 2.85E-52 | 1.490433 | 0.863 | 0.859 | 6.33E-48 |
| CFL2 | 3.10E-41 | 1.4873 | 0.543 | 0.32 | 6.87E-37 |
| ID3 | 3.36E-21 | 1.467823 | 0.474 | 0.323 | 7.45E-17 |
| EIF4A2 | 1.41E-92 | 1.453702 | 0.866 | 0.759 | 3.13E-88 |
| HMGB2 | 2.07E-39 | 1.440726 | 0.484 | 0.223 | 4.58E-35 |
| RPS3 | 7.07E-158 | 1.437167 | 0.995 | 0.936 | 1.57E-153 |
| RPS20 | 2.22E-134 | 1.436909 | 0.972 | 0.784 | 4.92E-130 |
| TPT1 | 1.25E-147 | 1.432744 | 0.998 | 0.957 | 2.78E-143 |
| OLFML3 | 4.91E-36 | 1.412725 | 0.662 | 0.559 | 1.09E-31 |
| RPS13 | 1.88E-142 | 1.410261 | 0.989 | 0.85 | 4.16E-138 |
| RPL9 | 4.10E-150 | 1.404303 | 0.989 | 0.893 | 9.10E-146 |
| TSC22D1 | 2.78E-31 | 1.403521 | 0.573 | 0.42 | 6.17E-27 |
| RPL37A | 3.41E-112 | 1.401321 | 0.935 | 0.73 | 7.57E-108 |
| RPS7 | 2.08E-151 | 1.399044 | 0.99 | 0.907 | 4.60E-147 |
| RPS15 | 1.23E-150 | 1.395557 | 0.996 | 0.916 | 2.73E-146 |
| RPL32 | 2.78E-142 | 1.394387 | 0.998 | 0.925 | 6.17E-138 |
| LSP1 | 1.92E-33 | 1.390285 | 0.342 | 0.08 | 4.25E-29 |
| SERPINE2 | 1.11E-44 | 1.38508 | 0.581 | 0.291 | 2.46E-40 |
| RPL35A | 1.75E-140 | 1.374752 | 0.99 | 0.855 | 3.87E-136 |

**Supplementary Table 8: Top significant DE genes comparing fetal and maternal fibroblast type 2 cells in placenta tissue, as identified by Originator. log2 fold change (avg_log2FC) is averaged expression in fetal compared to maternal fibroblast type 2.**

|  | **p_val** | **avg_log2FC** | **pct.1** | **pct.2** | **p_val_adj** |
| --- | --- | --- | --- | --- | --- |
| EGFL6 | 1.15E-97 | 2.618358 | 0.28 | 0.05 | 2.56E-93 |
| TCF21 | 2.16E-114 | 2.612331 | 0.271 | 0.032 | 4.79E-110 |
| GPC3 | 9.91E-104 | 2.512707 | 0.287 | 0.048 | 2.20E-99 |
| TFPI2 | 1.58E-46 | 2.028858 | 0.145 | 0.026 | 3.50E-42 |
| DLK1 | 2.05E-91 | 1.895907 | 0.273 | 0.051 | 4.55E-87 |
| COL3A1 | 1.31E-39 | 1.799687 | 0.442 | 0.273 | 2.91E-35 |
| MEG3 | 8.54E-56 | 1.785014 | 0.721 | 0.573 | 1.89E-51 |
| CD36 | 1.01E-93 | 1.757117 | 0.257 | 0.04 | 2.23E-89 |
| S100A10 | 6.75E-180 | 1.60851 | 0.782 | 0.354 | 1.50E-175 |
| HSD17B2 | 6.80E-73 | 1.600796 | 0.24 | 0.05 | 1.51E-68 |
| IGF2 | 4.09E-77 | 1.595697 | 0.885 | 0.745 | 9.07E-73 |
| IGFBP3 | 1.74E-132 | 1.491305 | 0.946 | 0.806 | 3.85E-128 |
| FN1 | 1.06E-108 | 1.485688 | 0.862 | 0.691 | 2.35E-104 |
| HAPLN1 | 2.43E-68 | 1.3866 | 0.128 | 0.005 | 5.39E-64 |
| PITX2 | 3.98E-119 | 1.384857 | 0.211 | 0.007 | 8.82E-115 |
| EPYC | 9.74E-99 | 1.382534 | 0.401 | 0.12 | 2.16E-94 |
| SERPINE2 | 1.19E-78 | 1.346266 | 0.445 | 0.18 | 2.64E-74 |
| SPARC | 2.66E-125 | 1.28373 | 0.964 | 0.811 | 5.89E-121 |
| PLA2G2A | 8.61E-23 | 1.26497 | 0.213 | 0.103 | 1.91E-18 |
| TAC3 | 1.77E-58 | 1.263857 | 0.444 | 0.213 | 3.92E-54 |
| WNT2 | 5.83E-64 | 1.259141 | 0.208 | 0.041 | 1.29E-59 |
| HGF | 3.71E-80 | 1.238979 | 0.174 | 0.015 | 8.23E-76 |
| PDPN | 4.69E-15 | 1.230947 | 0.549 | 0.486 | 1.04E-10 |
| C7 | 5.50E-51 | 1.230843 | 0.193 | 0.046 | 1.22E-46 |
| CYTL1 | 1.03E-53 | 1.219926 | 0.1 | 0.004 | 2.29E-49 |
| ENPP2 | 4.13E-50 | 1.171042 | 0.308 | 0.124 | 9.15E-46 |
| COL1A1 | 8.28E-32 | 1.14892 | 0.661 | 0.541 | 1.84E-27 |
| PHACTR2 | 1.05E-24 | 1.127201 | 0.369 | 0.248 | 2.32E-20 |
| CSH1 | 2.13E-29 | 1.125288 | 0.645 | 0.532 | 4.72E-25 |
| PRG2 | 4.21E-75 | 1.089949 | 0.572 | 0.325 | 9.33E-71 |
| RARRES2 | 1.68E-11 | 1.076482 | 0.23 | 0.153 | 3.72E-07 |
| IGFBP1 | 5.03E-13 | 1.076354 | 0.755 | 0.847 | 1.12E-08 |
| MATN2 | 4.80E-35 | 0.976729 | 0.23 | 0.093 | 1.06E-30 |
| TIMP2 | 2.72E-65 | 0.972391 | 0.819 | 0.783 | 6.02E-61 |
| PPDPF | 2.70E-126 | 0.966404 | 0.94 | 0.946 | 5.99E-122 |
| PMP22 | 2.37E-09 | 0.964084 | 0.422 | 0.382 | 5.26E-05 |
| COL1A2 | 2.51E-14 | 0.956048 | 0.417 | 0.333 | 5.56E-10 |
| GNAS | 4.56E-36 | 0.956025 | 0.727 | 0.673 | 1.01E-31 |
| TIMP3 | 1.26E-38 | 0.951015 | 0.929 | 0.969 | 2.78E-34 |
| TMEM176B | 5.65E-34 | 0.937679 | 0.564 | 0.365 | 1.25E-29 |
| KRT18 | 1.96E-23 | 0.934884 | 0.575 | 0.485 | 4.35E-19 |
| CD9 | 2.35E-63 | 0.926787 | 0.32 | 0.109 | 5.21E-59 |
| CNN3 | 1.70E-15 | 0.924418 | 0.345 | 0.248 | 3.76E-11 |
| TMEM98 | 7.11E-88 | 0.884145 | 0.821 | 0.705 | 1.58E-83 |
| TAGLN2 | 1.10E-32 | 0.868028 | 0.669 | 0.574 | 2.44E-28 |
| JAM2 | 7.44E-54 | 0.836138 | 0.573 | 0.396 | 1.65E-49 |
| ADAMTS1 | 8.64E-35 | 0.800778 | 0.761 | 0.686 | 1.91E-30 |
| KCNQ1OT1 | 3.23E-54 | 0.777925 | 0.587 | 0.378 | 7.15E-50 |
| CRLF1 | 5.44E-86 | 0.769451 | 0.361 | 0.106 | 1.21E-81 |
| CITED2 | 1.72E-22 | 0.737314 | 0.647 | 0.59 | 3.82E-18 |
| KRT8 | 2.11E-12 | 0.731913 | 0.391 | 0.312 | 4.67E-08 |
| CLU | 3.36E-38 | 0.730333 | 0.722 | 0.608 | 7.45E-34 |
| IL1RL1 | 2.00E-11 | 0.729334 | 0.453 | 0.369 | 4.43E-07 |
| C12orf39 | 2.18E-55 | 0.723933 | 0.106 | 0.005 | 4.84E-51 |
| RPS4Y1 | 2.55E-31 | 0.723712 | 0.212 | 0.083 | 5.66E-27 |
| CRISPLD2 | 4.10E-52 | 0.722449 | 0.778 | 0.676 | 9.08E-48 |
| VTN | 2.37E-14 | 0.720926 | 0.34 | 0.25 | 5.24E-10 |
| TGFBR1 | 3.40E-72 | 0.715112 | 0.385 | 0.146 | 7.54E-68 |
| ATP2B4 | 9.49E-33 | 0.708239 | 0.622 | 0.533 | 2.10E-28 |
| PAPPA | 1.11E-24 | 0.703654 | 0.619 | 0.533 | 2.46E-20 |
| PPIC | 4.13E-46 | 0.696404 | 0.686 | 0.52 | 9.17E-42 |
| TMEM176A | 3.69E-14 | 0.685517 | 0.435 | 0.318 | 8.18E-10 |
| TGFBI | 2.88E-16 | 0.678673 | 0.403 | 0.293 | 6.37E-12 |
| VEGFA | 1.51E-11 | 0.677592 | 0.409 | 0.326 | 3.35E-07 |
| PRUNE2 | 5.06E-45 | 0.677505 | 0.572 | 0.376 | 1.12E-40 |
| AOC1 | 1.09E-15 | 0.677382 | 0.357 | 0.252 | 2.41E-11 |
| PAPPA2 | 2.09E-17 | 0.673445 | 0.188 | 0.095 | 4.63E-13 |
| HTRA1 | 1.08E-28 | 0.672958 | 0.525 | 0.391 | 2.40E-24 |
| COL6A1 | 3.34E-13 | 0.671865 | 0.432 | 0.348 | 7.41E-09 |
| NOTUM | 2.95E-13 | 0.667101 | 0.198 | 0.116 | 6.54E-09 |
| CRABP2 | 3.17E-24 | 0.665274 | 0.174 | 0.07 | 7.03E-20 |
| TWISTNB | 2.47E-14 | 0.664275 | 0.697 | 0.697 | 5.48E-10 |
| OGN | 3.36E-32 | 0.663175 | 0.393 | 0.237 | 7.45E-28 |
| FXYD1 | 1.09E-50 | 0.663043 | 0.775 | 0.691 | 2.41E-46 |
| PARM1 | 2.92E-26 | 0.652733 | 0.649 | 0.572 | 6.47E-22 |
| IGFBP4 | 1.31E-11 | 0.64799 | 0.79 | 0.878 | 2.90E-07 |
| IGFBP2 | 2.75E-26 | 0.643782 | 0.768 | 0.757 | 6.10E-22 |
| WNT4 | 2.11E-18 | 0.636485 | 0.409 | 0.304 | 4.68E-14 |
| CYP26B1 | 1.89E-48 | 0.63562 | 0.249 | 0.082 | 4.20E-44 |
| LAMP1 | 8.50E-32 | 0.634004 | 0.462 | 0.313 | 1.88E-27 |
| VAMP8 | 3.27E-20 | 0.628365 | 0.302 | 0.191 | 7.25E-16 |
| FBN2 | 5.84E-34 | 0.624472 | 0.38 | 0.211 | 1.29E-29 |
| SLC25A4 | 6.00E-40 | 0.624015 | 0.547 | 0.365 | 1.33E-35 |
| LBH | 1.64E-57 | 0.621446 | 0.255 | 0.071 | 3.64E-53 |
| FBLN5 | 5.70E-36 | 0.618509 | 0.748 | 0.648 | 1.26E-31 |
| SERINC1 | 5.81E-46 | 0.61593 | 0.625 | 0.454 | 1.29E-41 |
| FBLN1 | 5.21E-48 | 0.612491 | 0.794 | 0.688 | 1.16E-43 |
| CD74 | 8.58E-14 | 0.61232 | 0.186 | 0.103 | 1.90E-09 |
| ALDH1A2 | 9.49E-27 | 0.60386 | 0.517 | 0.393 | 2.10E-22 |
| COX16 | 6.20E-35 | 0.60385 | 0.706 | 0.632 | 1.37E-30 |
| GSTA1 | 1.14E-34 | 0.595045 | 0.201 | 0.067 | 2.52E-30 |
| PGRMC2 | 1.59E-31 | 0.594757 | 0.461 | 0.304 | 3.53E-27 |
| VAMP2 | 4.78E-38 | 0.588629 | 0.713 | 0.655 | 1.06E-33 |
| CD248 | 8.25E-25 | 0.585625 | 0.776 | 0.778 | 1.83E-20 |
| NPC2 | 1.82E-61 | 0.583092 | 0.96 | 0.933 | 4.04E-57 |
| LSP1 | 8.32E-26 | 0.575824 | 0.251 | 0.127 | 1.84E-21 |
| NFE2L1 | 1.79E-26 | 0.568322 | 0.544 | 0.43 | 3.96E-22 |
| QSOX1 | 2.03E-19 | 0.56532 | 0.445 | 0.334 | 4.51E-15 |
| TNFRSF21 | 9.14E-24 | 0.559502 | 0.404 | 0.274 | 2.03E-19 |
| MALAT1 | 2.61E-95 | 0.557937 | 0.995 | 0.995 | 5.79E-91 |

**Supplementary Table 9: Top significant DE genes comparing fetal and maternal macrophages in placenta tissue, as identified by Originator. log2 fold change (avg_log2FC) is averaged expression in fetal compared to maternal macrophages.**

|  | **p_val** | **avg_log2FC** | **pct.1** | **pct.2** | **p_val_adj** |
| --- | --- | --- | --- | --- | --- |
| SEPP1 | 7.59E-10 | 2.939086 | 0.863 | 0.296 | 1.68E-05 |
| ANKRD22 | 1.82E-07 | -0.31847 | 0.005 | 0.185 | 0.004043 |
| HLA-DOA | 5.87E-09 | -0.41583 | 0.005 | 0.222 | 0.00013 |
| FBXO6 | 3.80E-08 | -0.50193 | 0.016 | 0.259 | 0.000842 |
| CD38 | 6.69E-09 | -0.51085 | 0.005 | 0.222 | 0.000148 |
| SIGLEC10 | 1.39E-10 | -0.51302 | 0.005 | 0.259 | 3.08E-06 |
| FGR | 1.85E-09 | -0.55798 | 0.016 | 0.296 | 4.11E-05 |
| TAGAP | 3.68E-09 | -0.55826 | 0.027 | 0.333 | 8.17E-05 |
| SLAMF7 | 2.07E-07 | -0.57248 | 0.005 | 0.185 | 0.004591 |
| APOBEC3A | 1.76E-07 | -0.63216 | 0 | 0.148 | 0.003903 |
| PILRA | 1.11E-07 | -0.68662 | 0.066 | 0.407 | 0.002469 |
| AC147651.4 | 5.15E-09 | -0.71519 | 0.005 | 0.222 | 0.000114 |
| CLEC4E | 1.27E-10 | -0.73408 | 0 | 0.222 | 2.82E-06 |
| SSPN | 9.46E-08 | -0.7427 | 0.038 | 0.333 | 0.002096 |
| FCN1 | 5.17E-09 | -0.78535 | 0.011 | 0.259 | 0.000115 |
| ENO1 | 2.82E-07 | -0.91871 | 0.835 | 0.963 | 0.00624 |
| GBP1 | 2.32E-08 | -1.01877 | 0.088 | 0.481 | 0.000515 |
| FBP1 | 8.28E-09 | -1.04538 | 0.066 | 0.444 | 0.000184 |
| GBP2 | 9.71E-10 | -1.06374 | 0.066 | 0.481 | 2.15E-05 |
| CD48 | 4.85E-11 | -1.07269 | 0.066 | 0.519 | 1.08E-06 |
| HLA-DQA2 | 7.92E-14 | -1.09034 | 0.016 | 0.407 | 1.76E-09 |
| TYMP | 8.40E-10 | -1.10807 | 0.044 | 0.407 | 1.86E-05 |
| STAT1 | 7.60E-10 | -1.15925 | 0.071 | 0.481 | 1.69E-05 |
| TNFSF13B | 7.24E-08 | -1.27052 | 0.154 | 0.593 | 0.001605 |
| GCHFR | 2.23E-08 | -1.27756 | 0.242 | 0.778 | 0.000494 |
| HLA-A | 1.32E-07 | -1.29365 | 0.775 | 1 | 0.002918 |
| HLA-DMA | 2.39E-07 | -1.2983 | 0.302 | 0.741 | 0.005308 |
| FCGR3A | 3.41E-07 | -1.36724 | 0.385 | 0.852 | 0.007564 |
| LSP1 | 1.88E-10 | -1.38666 | 0.148 | 0.704 | 4.16E-06 |
| CXCL9 | 3.32E-12 | -1.4032 | 0 | 0.259 | 7.35E-08 |
| LAPTM5 | 7.70E-08 | -1.44247 | 0.522 | 0.963 | 0.001706 |
| CAPG | 3.11E-08 | -1.46339 | 0.324 | 0.852 | 0.00069 |
| HAMP | 6.41E-10 | -1.5656 | 0.033 | 0.37 | 1.42E-05 |
| SOD2 | 3.87E-07 | -1.66751 | 0.346 | 0.778 | 0.008586 |
| PLAUR | 4.33E-08 | -1.6812 | 0.225 | 0.704 | 0.00096 |
| TREM2 | 7.34E-11 | -1.8716 | 0.176 | 0.704 | 1.63E-06 |
| CXCR4 | 6.03E-13 | -1.99228 | 0.066 | 0.556 | 1.34E-08 |
| HLA-DRB1 | 7.34E-12 | -2.13116 | 0.39 | 0.963 | 1.63E-07 |
| CD52 | 4.03E-10 | -2.16276 | 0.082 | 0.519 | 8.93E-06 |
| APOC1 | 1.80E-10 | -2.28506 | 0.258 | 0.815 | 4.00E-06 |
| HLA-DQB1 | 3.08E-15 | -2.28714 | 0.154 | 0.815 | 6.82E-11 |
| CXCL10 | 2.56E-09 | -2.34346 | 0.011 | 0.259 | 5.66E-05 |
| PLIN2 | 4.16E-10 | -2.37356 | 0.363 | 0.889 | 9.22E-06 |
| CD74 | 3.73E-11 | -2.48105 | 0.632 | 1 | 8.26E-07 |
| HLA-DRB5 | 1.19E-07 | -2.52589 | 0.198 | 0.593 | 0.002643 |
| LYZ | 2.09E-11 | -2.59556 | 0.264 | 0.815 | 4.64E-07 |
| HLA-DPB1 | 1.63E-13 | -2.61009 | 0.247 | 0.889 | 3.61E-09 |
| RGS1 | 4.57E-18 | -2.61249 | 0.077 | 0.704 | 1.01E-13 |
| HLA-DQA1 | 1.03E-12 | -2.68723 | 0.11 | 0.63 | 2.27E-08 |
| HLA-DRA | 1.38E-12 | -2.86693 | 0.462 | 0.963 | 3.06E-08 |
| HLA-DPA1 | 9.05E-16 | -2.89269 | 0.28 | 0.963 | 2.01E-11 |

**Supplementary Table 10: Cell-type-specific marker genes for placenta tissues**

| **cell type** | **genes** |
| --- | --- |
| Villous cytotrophoblast | PAGE4, PEG10, ISYNA1 |
| Syncytiotrophoblast | CGA, CYP19A1 |
| Extravillous trophoblast | HLA-G, HTRA4 |
| Fibroblast type 1 | DLK1, EGFL6, GPC3 |
| Fibroblast type 2 | DKK1, IGFBP5, IGFBP1 |
| Vascular endothelial cell | CD34, VWF, CLEC14A, ECSCR |
| T-cell | IL7R, CCR7 |
| NK cell | GNLY, NKG7 |
| Macrophage (HB) | LYVE1, DAB2, CCL4, CSF1R, CD163, CD209 |
| Monocyte | FCGR3A, MS4A7, CD14, LYZ |
| Erythrocyte | HBB, HBG1, HBA1 |

**Supplementary Table 11: Cell-type-specific marker genes for PDAC tissues**

| **cell type** | **genes** |
| --- | --- |
| Epithelial cell | KRT19 |
| Ductal cell | SLC3A1, VTCN1, DCDC2, SERPINA5 |
| Acinar cell | PRSS3, PNLIP, CTRC |
| Endothelial cell | CD34, VWF, KDR |
| Peri-islet Schwann cell | GFRA3, MPZ, GFRA1, INSC, SOX10, S100B |
| Beta cell | INS, INS2, MAFA |
| Fibroblast | COL1A1, ACTA2 |
| M2 macrophage | CD163, PPARG, MRC1 |
| Myeloid-derived suppressor cell (MDSC) | S100A9, ICAM1, S100A8, CXCR1 |
| T-cell | IL7R, CCR7 |
| Regulatory T-cell | IKZF2, FOXP3, CTLA4 |
| NK cell | GNLY, NKG7, NCR1 |
| B-cell | MS4A1, CD19, BLK |
| Mast cell | KIT |
| Plasma cell | MZB1, SPAG4 |

**Supplementary Table 12: Biological interpretation of exclusive genes between 1) T-cell and T-reg, 2) macrophage and NK cell, and 3) NK cell, T-cell, and macrophage**

| **Common cell types** | **Cell type** | **Gene** | **Highly express in** | | **Supporting citation** |
| --- | --- | --- | --- | --- | --- |
|  |  |  | **blood** | **Expected tissue-resident** |  |
| T- cell & T-reg | T-cell | MT-ND1 |  | ✓ | Lin et al. [1]; Rivadeinera & Delgoffe [2] |
|  |  | TNFRSF4 | ✓ |  | Ma et al. [3]; Iriki et al. [4]; Luo et al. [5] |
|  |  | RPS26 | ✓ |  | Chen et al. [6] |
|  |  | LTB | ✓ |  | Abdulrahman et al. [7] |
|  | T-reg | MT-ND1 |  | ✓ | Field et al. [8] |
|  |  | TNFRSF4 | ✓ |  | Chen et al. [9] |
|  |  | RPS26 | ✓ |  | Chen et al. [6] |
|  |  | LTB | ✓ |  | Abdulrahman et al. [7] |
| Macrophage & NK cell | Macrophage | RGCC |  | ✓ | Kamata & Tada [10]; Xu et al. [11] |
|  |  | CD63 | ✓ |  | Ye et al. [12]; Zhong et al. [13]; Khushman et al. [14] |
|  |  | LGALS1 | ✓ |  | Min et al. [15]; Murphy et al [16]; Schmieder & Schledzewski [17] |
|  | NK cell | RGCC |  | ✓ | Yu et al. [18] |
|  |  | CD63 | ✓ |  | Jewett et al. [19]; Khushman et al. [14] |
|  |  | LGALS1 | ✓ |  | Baker et al. [20]; Yu et al. [21] |
| NK cell, T-cell, & macrophage | NK cell | ZNF331 |  | ✓ | Marquardt et al. [22]; Foroutan et al. [23] |
|  | T-cell | ZNF331 | ✓ |  | Egelston et al. [24] |
|  | macrophage | ZNF331 | ✓ |  | Xiao et al. [25] |

25.	Xiao F, Shen J, Zhou L, Fang Z, Weng Y, Zhang C, et al. ZNF395 facilitates macrophage polarization and impacts the prognosis of glioma.
